# How striatal neurons influence decision-making depends on broader circuit context: evidence from theory-driven cell type identification

**DOI:** 10.64898/2026.11.10.765679

**Authors:** Zhuojun Yu, Timothy Verstynen, Jonathan E. Rubin

## Abstract

Neural cell types are often conceived as having fixed computational roles, yet their behavioral contributions may depend on the broader circuit state. In the striatum, for example, direct and indirect pathway spiny projection neurons (SPNs) are typically described as facilitating or suppressing actions, respectively. However, emerging observations of striatal activity, such as modulation related to action timing, action vigor, and reactive inhibition, call this dichotomy into question. Here we develop a state-space framework linking striatal population activity to decision dynamics by combining a spiking cortico-basal ganglia-thalamic (CBGT) model with large-scale *in vivo* recordings from mice performing a two-choice task. Modeled activity patterns guided the inference of putative striatal SPN subtypes in the empirical recordings, enabling reconstruction of decision trajectories through an underlying state space. Across simulated and experimental data, these trajectories revealed a shared set of decision dynamics and identified three state-dependent roles of indirect pathway activity: preserving uncertainty when active alone, signifying choice commitment when coactive with the direct pathway in the same action channel, and slowing decisions when recruited in the opposing channel. To interpret these roles mechanistically, we mapped state transitions onto previously described basal ganglia control ensembles (responsiveness, pliancy, and choice) that translate circuit activity into the evidence accumulation process, and we found that each role was associated with a distinct pattern of control ensemble engagement that tunes decision policy. These analyses support a context-dependent view of SPN function, in which their moment-to-moment impact emerges from the evolving profile of dynamics throughout the CBGT circuit.

**Significance:** Mammals adaptively adjust their decision policies in changing environments. The basal ganglia are brain circuits central to this process, but their pathways are often described simply as opposing routes that either promote or suppress action selection. By using computational modeling results to infer identities of neurons within large-scale brain recordings and analyzing both simulated and recorded data, this work shows that the same pathway can play different roles depending on the broader activity state of the circuit. This state-context view provides a flexible account of how striatal neural populations contribute to behavior, including the extent of deliberation that takes place before commitment to an action occurs.

---

Theories about the function of the basal ganglia and its constituent nuclei have been debated at least since Willis first proposed that the striatum serves as a node for sensorimotor functions (1). Over time, this interconnected system of subcortical regions, particularly the segments that form recurrent loops with the neocortex and thalamus (2, 3), has been linked with action gating (4, 5), action kinematics (6), skill learning (7, 8), language processing (9, 10), and more (11). Many of these putative roles represent largely speculative hypotheses based on phenomenological associations or clinical observations. More recent circuit-level theories have focused on how basal ganglia pathways are ideally structured for evidence accumulation for decision-making (12), regulating the vigor of movement (13, 14), or predictive sequencing (15, 16). Each of these theories rests on the assumption that individual cell populations in the basal ganglia have static functional roles that contribute to broader network computations.

The canonical framework for conceptually organizing basal ganglia function proposes a division of labor between two structurally and functionally distinct pathways: the *direct* pathway, which facilitates action selection, and the *indirect* pathway, which suppresses action selection (3, 17, 18). Central to this framework is the assumption that these pathways are embedded within multiple parallel action *channels* (3–5), each corresponding to a candidate behavior and each containing direct- and indirect-pathway components. However, recent anatomical, electrophysiological, and computational evidence indicates that the two pathways do not act simply as competing Go/No-Go routes (19– 22). Instead, a growing body of work supports a more dynamic and heterogeneous view of pathway interactions during decision-making (23–27). For example, indirect pathway activity can accompany action initiation when coactive with the direct pathway (28), contribute to lateralized movement control (29), and support monitoring and updating of action selection (30). These various observations imply that indirect pathway activity can be engaged across distinct behavioral contexts with different functional demands. This suggests that the roles of indirect pathway spiny projection neurons (SPNs) within the decision-making process might not be interpretable from cell identity alone, but also depends on both broader network and task contexts.

Here we set out to determine whether the activity of major striatal neuron populations reflects static functional roles throughout the period leading up to a decision or, alternatively, whether their contribution may evolve throughout the decision process, depending on the current network state. To this end, we combined a spiking computational model of the cortico-basal ganglia-thalamic (CBGT) circuit (Fig. 1A; (33, 34)) with previously collected and publicly available large-scale Neuropixels recordings from the striatum in mice (Fig. 1B; (31, 32)) as both performed a simple two-choice task. We used the model-derived reference patterns to assign putative striatal cell subtypes in empirical recordings and then reconstructed decision trajectories through a set of network states using a recently developed analytical state-space approach (34). This analysis recasts striatal activity in terms of evolving decision trajectories, allowing us to understand how pathway-specific activity supports different behavioral roles across distinct states that may arise within the decision process.

**Fig 1.**
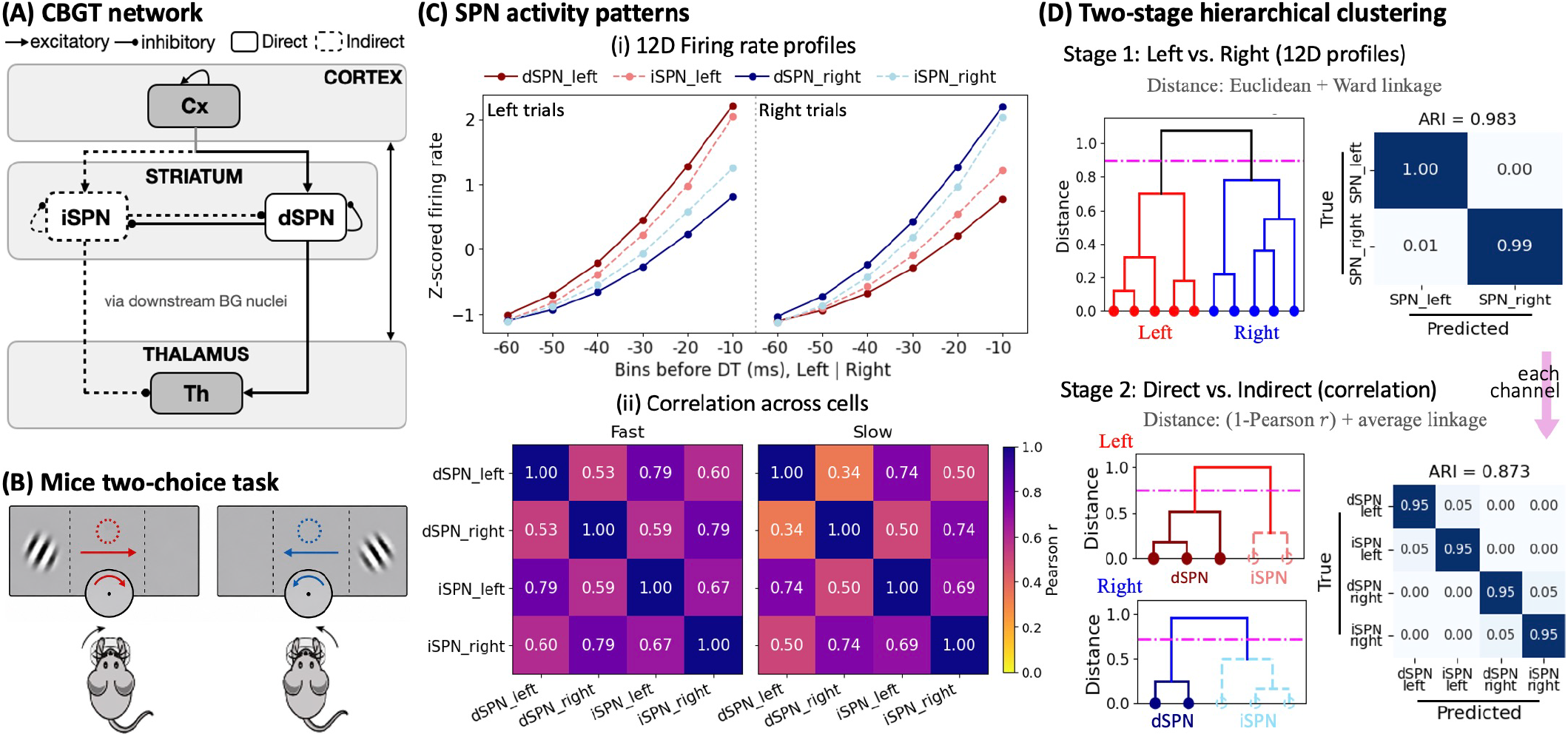
Model-guided inference of striatal state structure. (A) Reduced cortico-basal ganglia-thalamic (CBGT) circuit schematic showing the direct and indirect striatal pathways. The full simulated circuit and an example firing rate time course are shown in *SI Appendix*, Fig. S1. Cx, cortical neurons; dSPN, direct spiny projection neurons; iSPN, indirect spiny projection neurons; Th, thalamus. (B) Schematic of the visual two-choice task. Mice reported their choice by turning a wheel to move the selected visual grating to the center of the screen, clockwise for a left choice and counterclockwise for a right choice. Schematic adapted from (31, 32). (C) Simulated SPN activity patterns used for subtype inference. Panel i shows the firing rate profiles prior to decision, computed from the final six 10-ms bins before decision time and averaged separately for left- and right-choice trials, together comprising a 12D feature vector. Panel ii shows trial-level correlations between SPN units, measured by Pearson *r*, across fast and slow decision classes. (D) Two-stage hierarchical clustering approach for assigning SPN subtype labels. Stage 1 used the 12D firing rate profiles to separate left- and right-channel units, yielding an adjusted Rand index (ARI) of 0.983. Stage 2 used correlation distance, defined as 1 − *r*, within each inferred channel to separate dSPNs and iSPNs. The tree diagrams illustrate the clustering steps, with pink dashed horizontal lines indicating the dendrogram cut thresholds used to define clusters at each stage. The confusion matrices show classification performance in the simulated data, for which the true subtype labels were known. The complete two-stage procedure recovered the four SPN subtypes with high accuracy (ARI = 0.873).

## Results

### Striatal cell classification approach

We sought to clarify striatal function during a left–right decision-making task by using a computational framework to predict decision-related patterns of basal ganglia activity and testing whether these predictions matched empirical recordings. To do so, we first need to classify each untagged striatal neuron by both its pathway (direct or indirect) and its action representations (left or right action channels), yielding four possible class identities. The direct pathway starts with D1-expressing spiny projection neurons (dSPNs; solid in Fig. 1A) in the striatum, whereas the indirect pathway begins with D2-expressing spiny projection neurons (iSPNs; dashed). To establish reference activity patterns for these cells, we used 300 independently simulated CBGT networks in the context of a virtual two-choice task, reproduced from our previous work (34). These simulations provided labeled activity patterns for four pathway- and channel-specific populations: left-channel dSPNs, left-channel iSPNs, right-channel dSPNs, and right-channel iSPNs. The goal was to apply the same feature construction to pre-movement empirical activity recorded in mice performing a visual two-choice task, in which mice reported whether a visual stimulus appeared on the left or right by turning a wheel (Fig. 1B; (31, 32)), which we report in the next section. In this task, we assigned trials to hypothetical action channels based on the selected stimulus.

To construct features that could be compared directly across the simulated and empirical datasets, we first treated each of the four SPN populations within each CBGT network as if it were a single recorded unit. We then used pre-decision firing rate activity to derive two complementary features for distinguishing SPN subtypes. The first feature captured each unit’s average firing profile prior to decision. For each simulation run, which contained 50 trials from four SPN units, we aligned trials to decision time (DT) and extracted firing rates from the final six 10-ms bins before the decision. For each unit, we computed mean activity separately across left- and right-choice trials and concatenated these values into a 12-dimensional (12D) feature vector (see *SI Appendix*, Supporting text for more details). This feature reliably distinguished left and right action channels (Fig. 1C, panel i): left-channel units (red traces) showed stronger pre-decision ramping on trials ending in left choices, whereas right-channel units (blue traces) showed stronger ramping on trials ending in right choices.

However, this average pre-decision firing profile did not resolve pathway identity, because dSPNs and iSPNs within the same action channel could exhibit similar ramping patterns (see the red and blue pairs in Fig. 1C, panel i). We therefore introduced a second feature based on trial-to-trial covariation between SPN units. The simulated networks were divided into fast, intermediate, and slow decision-speed classes according to their overall DT distributions (*SI Appendix*, Fig. S1C). Within each class, we computed pairwise correlations among the four SPN units using their pre-decision firing rates across trials (Pearson *r*). This feature captured how SPN units covaried across trials and how these covariation patterns changed with decision speed (Fig. 1C, panel ii). The resulting correlation structure revealed a pathway-sensitive signature: correlations between left- and right-channel dSPNs decreased substantially from fast to slow decisions (0.53→0.34), whereas correlations between left- and right-channel iSPNs remained relatively stable (0.67→0.69). Hence, this feature allowed us to distinguish direct-from indirect-pathway SPN units within each action channel.

Based on these two features, we developed a two-stage hierarchical clustering approach to assign labels to the four latent SPN subtypes in any dataset (Fig. 1D). Stage 1 used the 12D firing rate profiles to separate left-channel and right-channel units. The labels inferred for the units in our simulations closely matched the known channel identities, achieving an adjusted Rand index (ARI) of 0.983. A forward bin-inclusion analysis also showed that channel recovery was dramatically stronger when all six pre-decision bins from each choice condition were included than when the feature was constructed from fewer bins (*SI Appendix*, Fig. S2), supporting the use of the complete 12D representation.

In Stage 2, units within each inferred channel were further separated into dSPNs and iSPNs using trial-level correlation structure, with dissimilarity quantified as *D* = 1 − *r*. In the simulated data, the resulting four-subtype classification agreed well with the known subtype identities (ARI = 0.873). Consistent with these high ARI values, the corresponding confusion matrices were strongly concentrated along the diagonal (Fig. 1D), indicating that most units were assigned to the correct action channel and SPN subtype.

### Striatal cell classification in recorded data

We next applied the same clustering procedure to two publicly available datasets, the Steinmetz dataset (31) and the International Brain Laboratory (IBL) dataset (32), with recordings of striatal populations during a two-choice visual task (Fig. 1B) in which mice reported their choice by turning a wheel in an option-specific direction. Because neither molecular pathway identity nor action-channel membership was directly measured in either dataset, we treated the inferred subtypes from the classifier as putative functional labels and evaluated their internal consistency and predictive utility in subsequent analyses. These evaluations used activity patterns, shuffle controls, and state-space analyses that were not themselves used to assign each label.

For each striatal unit, we constructed the same feature set used in the model, i.e., the 12D average firing rate profiles and trial-level correlation measures. Based on the reported motor-evoked potential latencies in mice, we defined the empirical decision-time proxy as the time point 10 ms before detected movement onset to minimize the influence of motor execution on the estimate (35, 36). For each dataset, we applied the filtering and quality-control criteria reported in its source publication. Before subtype assignment, we additionally excluded units with flat or decreasing pre-decision activity to focus our analysis on responsive units (*SI Appendix*, Supporting text).

Across sessions, the inferred action-channel groups showed choice-selective activity consistent with their assigned identities, and the inferred pathway groups exhibited temporal dynamics consistent with model predictions (example sessions in Fig. 2; all eligible sessions in *SI Appendix*, Figs. S3 and S4). First, units assigned to the left-action channel (red traces) ramped more strongly before rightward wheel reports, whereas units assigned to the right channel (blue traces) showed stronger ramps before leftward reports. Second, within each action channel, putative SPNs often showed similar pre-movement ramping, consistent with the model result that average firing profiles alone did not resolve pathway identity (compare Fig. 1C). Third, their trial-to-trial correlation patterns across decision-speed classes followed pathway-sensitive patterns predicted by the model (*SI Appendix*, Figs. S3 and S4), enabling recorded striatal units to be further separated into direct- and indirect-pathway subtypes. These inferred labels allowed us to link the empirical data to the theoretical model and investigate the functional contributions of pathway activation during decision-making.

**Fig 2.**
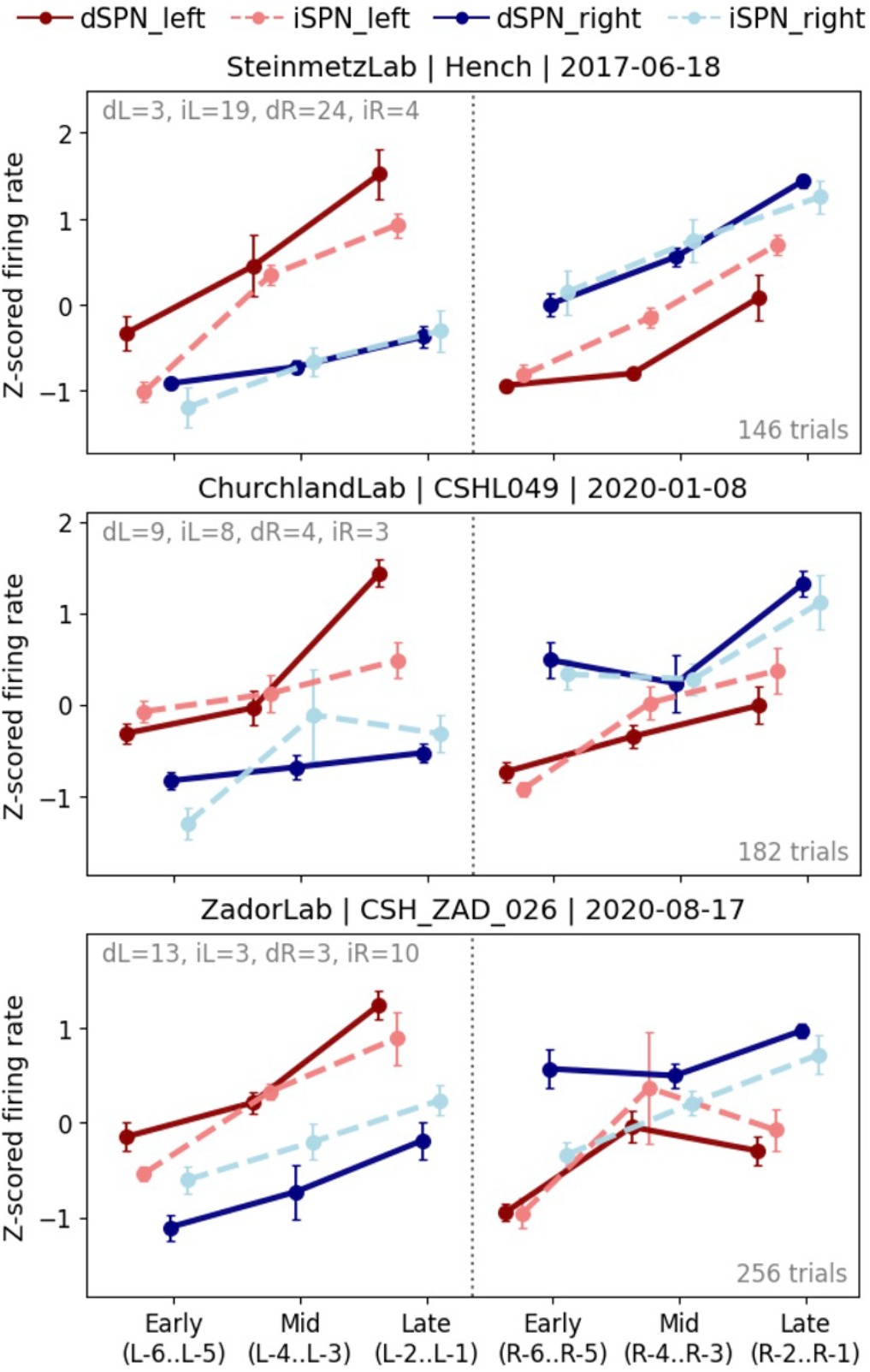
Putative SPN subtypes assigned to responsive striatal units in Neuropixels recordings (data from (31, 32)). Pre-movement firing rate profiles are shown for three example sessions (results for all eligible sessions are provided in *SI Appendix*, Figs. S3 and S4). Firing rates were z-scored and averaged across three pre-movement windows: early, middle, and late, corresponding to bins 6–5, 4–3, and 2–1 before decision. Colors denote inferred action channel, and line style denotes inferred pathway identity. Gray text reports the number of units assigned to each subtype, and trial counts are shown at the lower right of each panel. Error bars indicate SEM.

To determine whether this classification depended on specific temporal and task-related relationships, we applied three complementary shuffle controls, each designed to disrupt a distinct source of structure: spike timing, choice labels, or trial grouping by decision speed (*SI Appendix*, Fig. S5). The first shuffle control randomized the temporal ordering of each unit’s interspike intervals (ISIs) while preserving its spike count and ISI distribution. The choice-label shuffle reassigned left and right choices while preserving the number of trials in each group. The decision-speed shuffle reassigned fast and slow labels within each choice condition while preserving the corresponding group sizes. Across sessions, the subtype-specific temporal activity patterns in the original data were substantially more consistent with model predictions than those in the shuffled data (see *SI Appendix*, Fig. S5). Therefore, preserving only each unit’s overall firing statistics or the number of trials assigned to task labels was insufficient to reproduce the empirical patterns that supported the subtype assignments.

Finally, because the striatal recordings lacked genetic cell-type identification, we checked for the presence of fast-spiking interneurons (FSIs) in both empirical datasets. A unit was flagged as a putative FSI when its extracellular waveform was narrower than 0.15 ms at half of its maximum negative deflection and its whole-session firing rate exceeded 10 Hz (37–39). Among the clustered units, 9 of 164 Steinmetz units (5.5%) and 21 of 180 IBL units (11.7%) met both criteria (*SI Appendix*, Figs. S6 and S7). Importantly, excluding these units left the subtype-averaged activity profiles nearly unchanged, with pooled before-versus-after correlations of *r* = 0.998 for Steinmetz and *r* = 0.997 for IBL datasets, indicating that the subtype-specific activity patterns derived from the classification were not driven by the putative FSI units. We removed those putative FSI units and retained the remaining putative SPNs, together with their inferred subtype labels, in the subsequent state-space analyses.

### Decision trajectories in state space

We next characterized moment-by-moment decision trajectories through an underlying state space of unique network activity using a previously described CLAW (Circuit Logic Assessed via Walks) framework (34, 40). The CLAW method was adapted to a CBGT state space defined by activity of just four striatal populations. Within each time bin, firing rates from the four SPN groups were discretized, averaged, and binarized relative to a predefined threshold (*SI Appendix*, Fig. S1D). Striatal activity was then represented as a binary state vector, *s*_*k*_ = [dL, dR, iL, iR] ∈ {0, 1}^4^ (*k* = 0, 1, ⋯, 15), where each element of *s*_*k*_ is either 1 or 0, indicating whether the corresponding SPN group was active or not. In this way, each time bin was assigned to one of 16 possible states, and each continuous firing rate time series was converted into a sequence of discrete states, each representing the pattern of striatal activity in a time window. By aggregating state transitions across trials, we constructed high-probability transition chains, the *CLAW diagram*, which captured the dominant flow of striatal activity throughout the decision-making process (see (34, 40) and *SI Appendix*, Supporting text for details of the CLAW approach). We applied this method separately to the simulated CBGT activity with known SPN labels and to the empirical recordings with inferred SPN identities, enabling comparison of model and empirical decision trajectories within a common striatal state space (Fig. 3).

**Fig 3.**
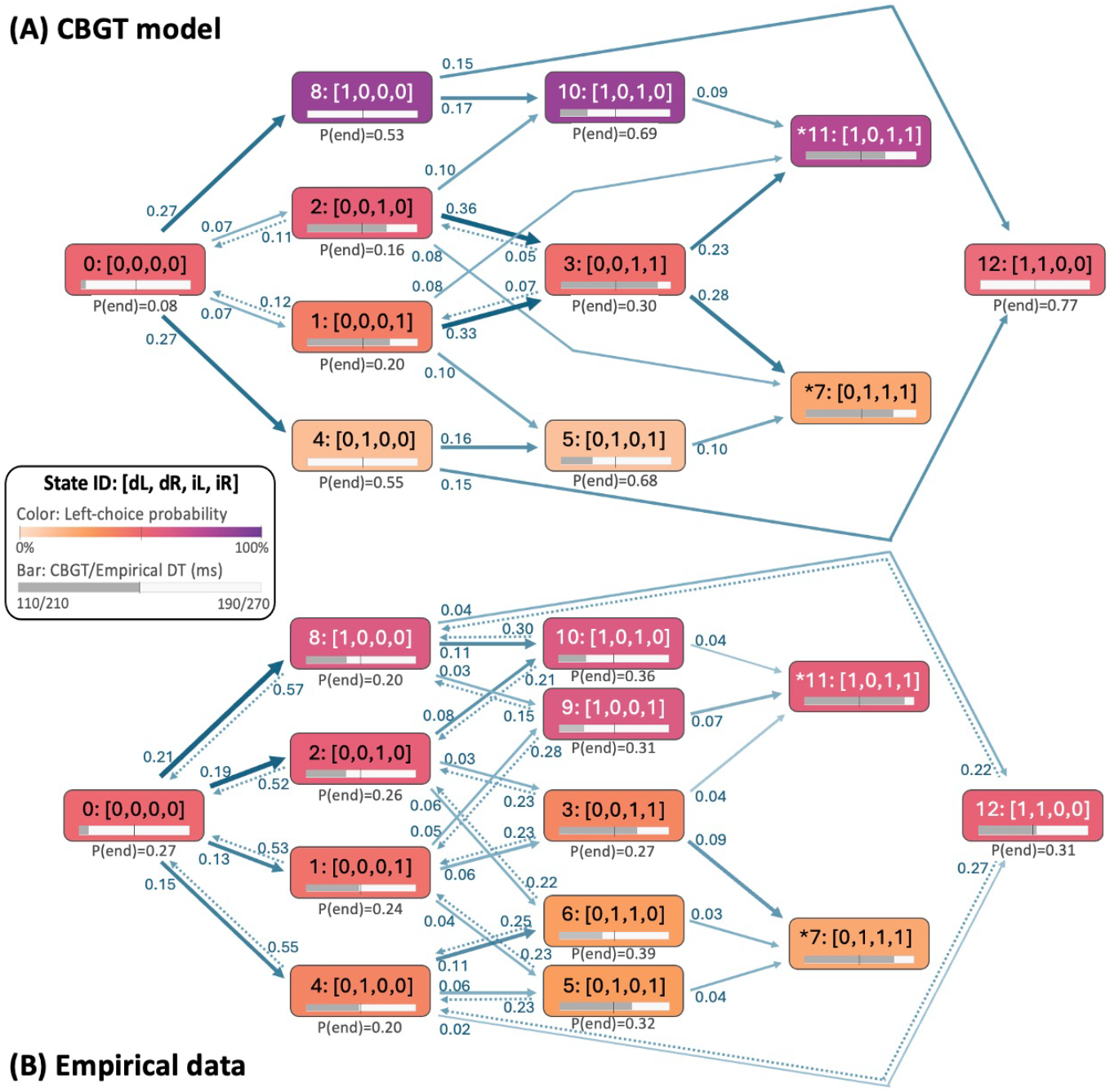
CLAW (Circuit Logic Assessed via Walks) diagrams for striatal dynamics in (A) simulated CBGT networks and (B) pooled empirical recordings from the IBL and Steinmetz datasets (31, 32), using the inferred SPN subtype labels. Each node represents a striatal state, labeled as state ID: [dL, dR, iL, iR], where dL and dR denote binarized left- and right-channel dSPN activity and iL and iR denote binarized left- and right-channel iSPN activity. A value of 1 indicates that the corresponding SPN group is active relative to a predefined threshold, whereas 0 indicates that it is inactive. Node color represents left-choice probability of trials visiting each state, ranging from low values in light orange (more right choices) to high values in purple (more left choices). The horizontal gray bar inside each node indicates the mean decision time of trials visiting this state, with separate ranges for the CBGT model and empirical data shown in the legend. The value below each node gives the terminal probability, *P* (end), defined as the probability that if this state is reached, then it is the final state prior to decisions. Arrows indicate transitions between states: solid arrows show forward transitions, with arrow width and color strength reflecting transition probability; dotted arrows show backward transitions. The number near each arrow gives the transition probability from the source state to the subsequent state. Asterisks mark low-frequency states 7 and 11; transitions from these states were not shown because they occurred at even lower probabilities.

In both the model and empirical CLAW diagrams, trajectories began from the inactive striatal state, [0, 0, 0, 0], that we denote as state 0, in which all SPN groups were below binarization threshold. This state served as a common *launching* state for subsequent decision dynamics. From there, activity either entered less choice-specific states in which the decision remained uncertain (inner CLAW) or branched into left- and right-biased paths (outer CLAW). States containing left-channel dSPN activation were associated with higher left-choice probability (see the node color of states 8, 10, and 11 in Fig. 3), whereas states containing right-channel dSPN activation were associated with lower left-choice probability (states 4, 5, and 7). This separation indicates that early activation of dSPNs in a specific channel marked the emergence of a choice-biased trajectory toward the corresponding action. In contrast, inner states 1, 2, and 3, which contained only iSPN activation, showed a weaker choice bias, indicating that iSPN-dominated activity was associated with uncertain states in which the two options were still being weighed. The empirical CLAW, based on real cell data (Fig. 3B), closely matched the choice structure predicted by the model (Fig. 3A), preserving the distinction between inner uncertain states versus outer choice-biased states.

Both CLAW diagrams further showed that complex circuit dynamics could be functionally divided into two critical phases: *deliberation* and *commitment*. The inner CLAW states had relatively low terminal probability (*P* (end) shown at the bottom of each node in Fig. 3) and multiple possible outgoing transitions, indicating that the trajectory remained in a deliberative phase in which the decision was not yet settled and could still evolve toward different outcomes. By contrast, forward progression through the outer CLAW paths led to states with higher *P* (end), corresponding to a commitment phase where activity became increasingly aligned with one action channel and more likely to end in the corresponding choice. In particular, same-channel dSPN+iSPN coactivation states (states 5 and 10) were among the most frequent states immediately preceding a decision. This late recruitment of iSPN activity after dSPN activation within the same action channel suggests that indirect pathway activity did not simply suppress action; rather, in this state context, it was associated with the final narrowing of the decision trajectory toward the choice already favored by the circuit.

We also observed two additional state features shared by the model and empirical CLAWs. First, the presence of a *reversal* state [1, 1, 0, 0], labeled as state 12, defined by activation of both left- and right-channel dSPNs, suggested that competing action channels could still play an important role even late in the decision process. This state appeared only after trajectories had already entered an outer CLAW path through states 4 or 8. Instead of continuing forward along that path, some trials entered state 12, where choice probability became more balanced relative to the preceding outer states (compare the node color of state 12 versus states 4 and 8). This reduction in choice bias indicates that the initial favoritism toward an action for trajectories along the outer CLAW was strong but not absolute. In this case, delayed ramping in the competing direct pathway could override the earlier bias and rapidly redirect the decision at the last moment, providing a reversal mechanism through which direct- pathway competition reshaped the final stage of decision formation. Second, both CLAWs shared a late recruitment of opponent-channel iSPN activity after same-channel dSPN+iSPN coactivation (states 5→7 and 10→11). In these transitions, activity first reached a state with dSPN and iSPN coactivation within the initially favored channel, after which activation in the opposing channel occurred. Compared with states 5 and 10, states 7 and 11 showed longer decision times (gray bars), indicating that late engagement of opponent-channel iSPN reflected rising competition from the opposing channel, which weakened the influence of the initially dominant channel and slowed the approach to action execution.

Alongside these shared features, the empirical CLAW was more variable than the model CLAW, with the addition of states 6 and 9, more frequent back-and-forth transitions between states, and less distinct separation of decision times between the inner and outer CLAWs. These differences likely reflect both biological variability and measurement constraints: empirical SPN identities were inferred from extracellular recordings, movement onset served only as an approximate proxy for decision time, and neural dynamics varied across trials and sessions due to training experience (see Discussion). Nevertheless, the core topology was preserved: a common launching state, branching left- and right-channel paths, inner deliberative states with iSPN dominance versus outer commitment states with dSPN dominance, the existence of a reversal state, and forward progression toward states with higher terminal probability. This agreement indicates that the inferred SPN subtypes captured a meaningful empirical state space and allowed the activity dynamics of individual trials to be represented as flows through that space.

### Context-dependent roles of iSPN activity

The CLAW comparisons led to three novel predictions about the state-dependent roles of indirect-pathway activity. First, isolated iSPN activation early in the trajectory characterized the deliberative states (states 1, 2, and 3), suggesting that it helps maintain *choice uncertainty* before either action channel dominates. Second, coactivation of dSPNs and iSPNs within the same channel occurred along the outer paths close to the time of action selection, suggesting that this configuration marks increasing *commitment* toward the corresponding choice (states 4→5 and 8→10). Third, subsequent recruitment of opponent-channel iSPNs (states 5→7 and 10→11) was associated with *delayed action selection*, consistent with increased competition from the opposing channel that delays resolution. These predictions define a state-dependent account in which the functional role of the indirect pathway depends on the broader striatal population state, rather than conforming to a fixed Go/No-Go rule.

We evaluated whether these predictions held across both simulated CBGT and empirical trials (Fig. 4). For each prediction, we identified the relevant CLAW states or transitions and quantified the behavioral measure most directly related to the proposed role: left-choice probability as a measure of uncertainty (panel A), terminal probability as a measure of commitment (panel B), and decision time as a measure of delayed action selection (panel C). Probability measures were assessed using trial-level bootstrap distributions, while decision-time effects were tested using the raw trial distributions (Materials and Methods). Simulation results and empirical data were analyzed separately.

**Fig 4.**
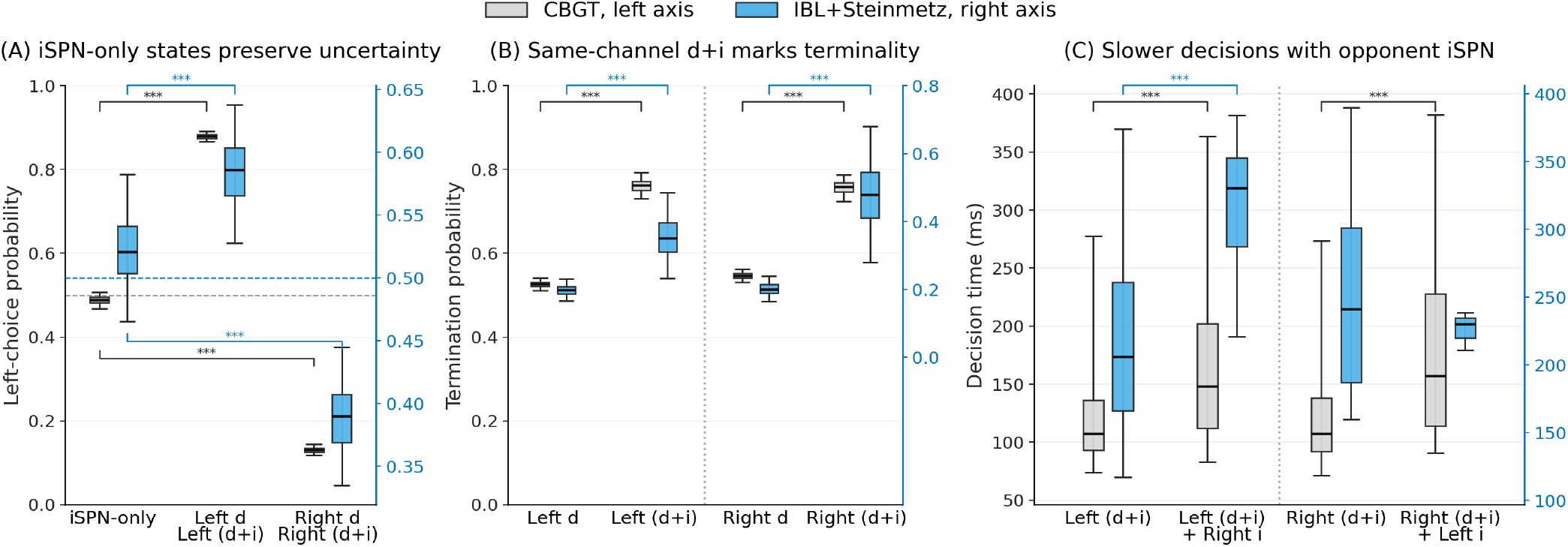
Model and empirical tests of the context-dependent roles of iSPN activity. Gray boxplots show CBGT model results using the left y-axis; blue boxplots show empirical results across recordings (31, 32) using the right y-axis. The two y-axes in each panel were scaled and aligned to facilitate visual comparison of the direction and consistency of effects between the model and empirical data. Panels A and B represent bootstrap distributions, and panel C represents raw trial-level decision time distributions. Boxes show the central 50% of trials or bootstrap samples, whiskers show the central 95%, and horizontal lines indicate medians. Black significance brackets correspond to CBGT model comparisons, and blue brackets correspond to empirical comparisons. Significance levels are indicated as \*\*\**p <* 0.001. (A) iSPN-only states preserve uncertainty. Left-choice probability was compared across iSPN-only states, left-channel dSPN states with or without same-channel iSPN coactivation (Left d, Left (d+i)), and right-channel dSPN states with or without same-channel iSPN coactivation (Right d, Right (d+i)). In both simulated and empirical data, the left choice probability for iSPN-only states remained near chance (0.5; horizontal dotted lines), whereas the left-channel dSPN states were biased toward left choices and the right-channel dSPN states were biased away from them. (B) Same-channel dSPN+iSPN coactivation marks terminality. For the left path, terminal probability was compared between all visits to state 8, labeled “Left d”, and visits to state 10, labeled “Left (d+i)”, that were entered directly from state 8. The corresponding right-path comparison contrasted all visits to state 4 (“Right d”) with visits to state 5 (“Right (d+i)”) that were entered directly from state 4. In both paths, adding same-channel iSPN activity was associated with higher terminal probability. (C) Slower decisions with opponent iSPN. Among trials that visited state 10 (“Left (d+i)”), decision times were compared according to whether the trajectory subsequently entered state 11 (“Left (d+i) + Right i”). The corresponding right-path analysis compared trials that visited state 5 (“Right (d+i)”) according to whether they subsequently entered state 7 (“Right (d+i) + Left i”). In the model, opponent-channel iSPN recruitment was associated with significantly longer decision times in both transitions; in the empirical data, this association was significant along the left path.

To test the first prediction, we compared left-choice probability among trials visiting iSPN-only states and those visiting dSPN-containing states on either the left or right path of the CLAW (Fig. 4A). States 1, 2, or 3, grouped as “iSPN-only”, occupied the inner region of the CLAW and contained iSPN activation without concurrent dSPN activation. In both simulated and empirical trials, visits to these states were associated with left-choice probabilities near chance (leftmost boxplots near the dotted horizontal lines). By contrast, states 8 and 10, pooled into the left-channel group (labeled “Left d” and “Left (d+i)” in the panel), were biased toward left choices; states 4 and 5, pooled into the right-channel group (“Right d” and “Right (d+i)”), were biased toward right choices. These biases support the interpretation that iSPN-only activity characterizes a regime of deliberation and uncertainty before dSPN engagement establishes bias toward a particular action channel.

The relationship changed when iSPNs were recruited within an action channel that already contained active dSPNs. Our model predicts that direct 8→10 transitions on the left path and 4→5 on the right path, which added same-channel iSPN activity, should be associated with trajectory termination (i.e., choice times). Indeed, when reached through these transitions, the coactivation states achieved significantly higher terminal probabilities than the preceding dSPN-only states in both simulated and empirical trials (Fig. 4B). Thus, once a choice-biased trajectory had already formed, iSPN activation in the same channel was associated with consolidation of selection rather than a return to uncertainty. This result argues against the traditional view that iSPN activity simply opposes action selection.

Finally, the model predicts that when iSPNs are subsequently recruited in the opposing action channel, decision times should be slower. On left-biased trajec-tories, trials that reached state 10 (labeled “Left (d+i)” in Fig. 4C) were separated according to whether they ended in decision or entered state 11 (“Left (d+i) + Right i” in Fig. 4C), which added right-channel iSPN activation to the existing left-channel dSPN+iSPN state. The corresponding right path examined whether trials reaching state 5 (“Right (d+i)”) subsequently entered state 7 (“Right (d+i) + Left i”) through the addition of left-channel iSPN activation. Because all compared trials had first reached the same-channel coactivation state, differences in decision time were specifically associated with the later emergence of iSPN activation in the opposing channel. In the CBGT model, this activation was associated with significantly longer decision times along both paths, and the empirical data reproduced the effect along the left path, reflecting an association between opponent-channel iSPN engagement and delayed action selection.

These state-level analyses establish that the behavioral roles of iSPN activity changed with its broader striatal context. In the final section, we analyzed the circuit mechanisms underlying these different roles and their consequences for decision-related policy.

### Circuit mechanisms underlying iSPN roles

The context-dependent role of iSPN activation supports the idea that the computational architecture of CBGT pathways has a low-dimensional structure. Our previous work (27, 34, 41) has identified three low-dimensional CBGT subnetworks, termed *control ensembles*, that map distinct patterns of circuit activity onto parameters of a normative drift-diffusion model (DDM; see (42)), which describes decision-making as the accumulation of evidence toward a boundary. These control ensembles provide a mechanistic bridge between the activity of specific CBGT populations and the tuning of specific aspects of decision policy (Fig. 5, left subpanels). Here, we utilized this mapping to project activity flow through the striatal CLAW onto the control ensembles and the corresponding DDM parameters, allowing us to derive an algorithmic interpretation of how the state transitions underlying each iSPN role shaped the decision outcomes. The *pliancy* and *responsiveness* ensembles (panels A and B) are primarily driven by the overall CBGT activity summed across both channels. Pliancy loads most strongly on indirect pathway activity, whereas responsiveness arises mainly from corticothalamic and direct pathway activity. Both ensembles modulate boundary height *a* (the amount of evidence needed to make a decision) and shorten the onset time *t* (the time at which evidence accumulation begins), but they exert opposing effects on *a*: pliancy raises the boundary required for action selection, while responsiveness lowers it. Therefore, the activation level of the responsiveness ensemble directly maps to the ease of making a decision; in a more nuanced way, increasing activation of the pliancy ensemble corresponds to earlier evaluation of evidence but the need for a higher standard of evidence before commitment to an action (i.e., less pliancy). In contrast to pliancy and responsiveness, the *choice* ensemble (panel C) is defined primarily by activity differences between the left and right action channels across all CBGT populations. This subnetwork encodes the dominance of one action channel over the other and mainly modulates drift rate *v* (the speed and direction of evidence accumulation toward a decision).

**Fig 5.**
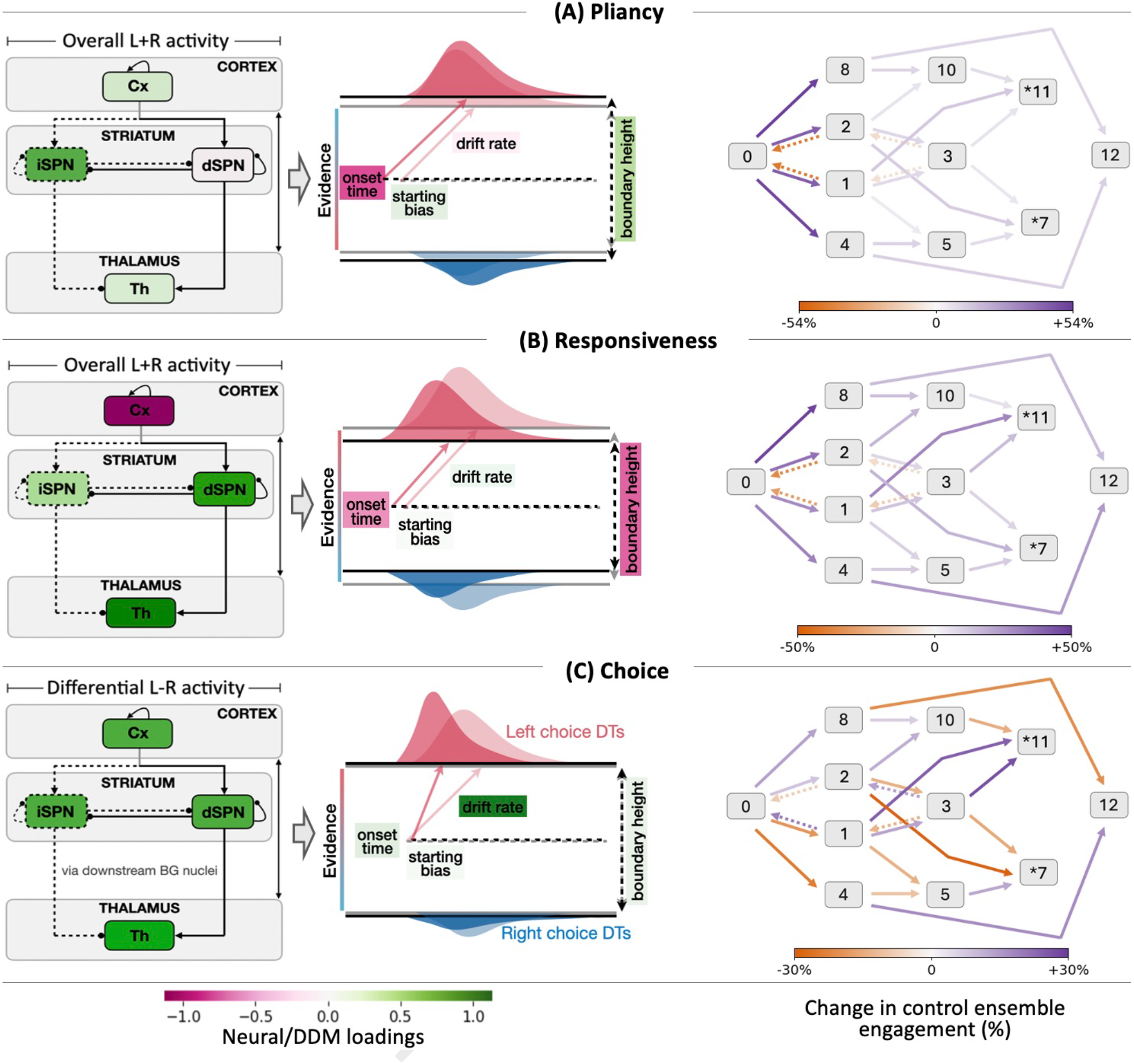
Control ensemble influence on striatal CLAW transitions. Panels A, B, and C represent the pliancy, responsiveness, and choice control ensembles, respectively, as manifested within the corticothalamic and striatal components of our CBGT model. Each left subpanel shows the relation between the firing rates of populations in the CBGT circuit and the DDM parameters, as identified by canonical correlation analysis (CCA). The control ensemble schematics are adapted from (40). The pliancy and responsiveness ensembles are associated with the overall activity summed across both channels (L+R); the choice ensemble is associated with the differential activity between left and right channels (L−R). The other aspect of activity for each ensemble (L–R for pliancy/responsiveness and L+R for choice) is not shown, because it contributed weakly to the corresponding DDM parameters. Green and pink shading indicate positive and negative canonical loadings, respectively. The accompanying DDM schematics depict how each control ensemble changes decision policy parameters and the resulting DT distributions. Pliancy and responsiveness both modulate boundary height and onset time, but exert opposing effects on boundary height. Choice primarily modulates drift rate. The CCA loadings for the full CBGT circuit are illustrated in *SI Appendix*, Fig. S8. Each right subpanel shows the projection of striatal CLAW transitions onto three control ensembles, using the same CLAW topology as Fig. 3A, with arrows colored according to the percentage change in control ensemble engagement from the preceding state to the subsequent state, relative to its engagement in the preceding state. Orange indicates decreased engagement, purple indicates increased engagement, and darker colors indicate changes of greater magnitude.

We projected each transition in the CBGT striatal CLAW onto the individual control ensembles to characterize how the engagement of CBGT subnetworks dynamically evolved as trajectories moved through the state space (Fig. 5, right subpanels) and how these changes shaped decision policy (*SI Appendix*, Fig. S9). We did not perform this analysis on the empirical CLAW because the control ensemble mapping requires activity measurements from the full set of CBGT populations (see Materials and Methods). The empirical recordings did not sample many of these populations simultaneously within individual sessions and did not provide the channel and subtype identities of certain recorded units (e.g., prototypical versus arkypallidal neurons in the external globus pallidus). Nevertheless, the agreement between model and empirical CLAWs in the three predictions (Fig. 4) allowed the model-based control ensemble analysis to provide mechanistic hypotheses for their shared state-dependent patterns.

Projecting the trajectories onto the control ensembles revealed distinct changes as they progressed through the CLAW. A clear distinction in activation of the choice ensemble emerged between entry into the left-versus right-biased outer paths (right subpanel of Fig. 5C), indicating that the corresponding L−R differential activity became increasingly positive along the left-biased path and increasingly negative along the right-biased path (i.e., greater R−L activity). Consequently, transitions toward left-channel dSPN states shifted the choice ensemble toward leftward drift, whereas transitions toward right-channel dSPN states produced the opposite shift (*SI Appendix*, Fig. S9B). These directional shifts in drift rate mirrored the choice biases associated with the corresponding CLAW states.

In contrast to the choice ensemble, pliancy and responsiveness both consistently increased as trajectories progressed forward through the CLAW (right subpanels of Fig. 5A and B), indicating growing engagement of the two CBGT subnetworks that regulate deliberative control and commitment readiness, respectively. Their joint increase is important because the two control ensembles have opposing effects on boundary height, where responsiveness promotes commitment by lowering the decision threshold but pliancy counterbalances this effect by raising the threshold. Thus, forward progression through the CLAW did not reflect a simple monotonic shift toward commitment driven by corticothalamic and direct pathway activity (activation of the responsiveness ensemble) but, instead, simultaneous engagement of the indirect pathway (activation of the pliancy ensemble) acted as a regulatory control that limited premature action selection.

Importantly, this counterbalancing effect did not prevent commitment. When the contributions of pliancy and responsiveness were combined and mapped onto boundary height, trajectories showed a consistent boundary collapse as they approached termination (*SI Appendix*, Fig. S9A). Thus, indirect pathway activity did not merely function as a brake on action selection; instead, it regulated the timing of commitment by prolonging deliberation, but as evidence accumulated and the decision trajectory advanced, the net policy still shifted toward a lower evidence threshold and eventual action selection.

These dynamic interactions of control ensembles provide a mechanistic account of the three state-dependent iSPN roles identified in Fig. 4. First, the uncertainty associated with iSPN-only activity in the inner CLAW was explained through the *pliancy* ensemble. Because pliancy is primarily driven by summed indirect pathway activity across channels and raises boundary height, iSPN activation early in the decision process strengthened pliancy, which sustained the evidence threshold and opposed premature commitment. Consistent with this interpretation, decision boundaries collapsed along the outer paths 0→8→10 and 0→4→5 but remained comparatively stable during transitions among states 1, 2 and 3 (*SI Appendix*, Fig. S9A). These inner iSPN-only states illustrate the counterbalancing role of pliancy relative to responsiveness: delaying the conversion of accumulated evidence into action so as to preserve deliberation. In Fig. 6, we schematically illustrate this role of iSPN-only activity along the route from launching to deliberation.

**Fig 6.**
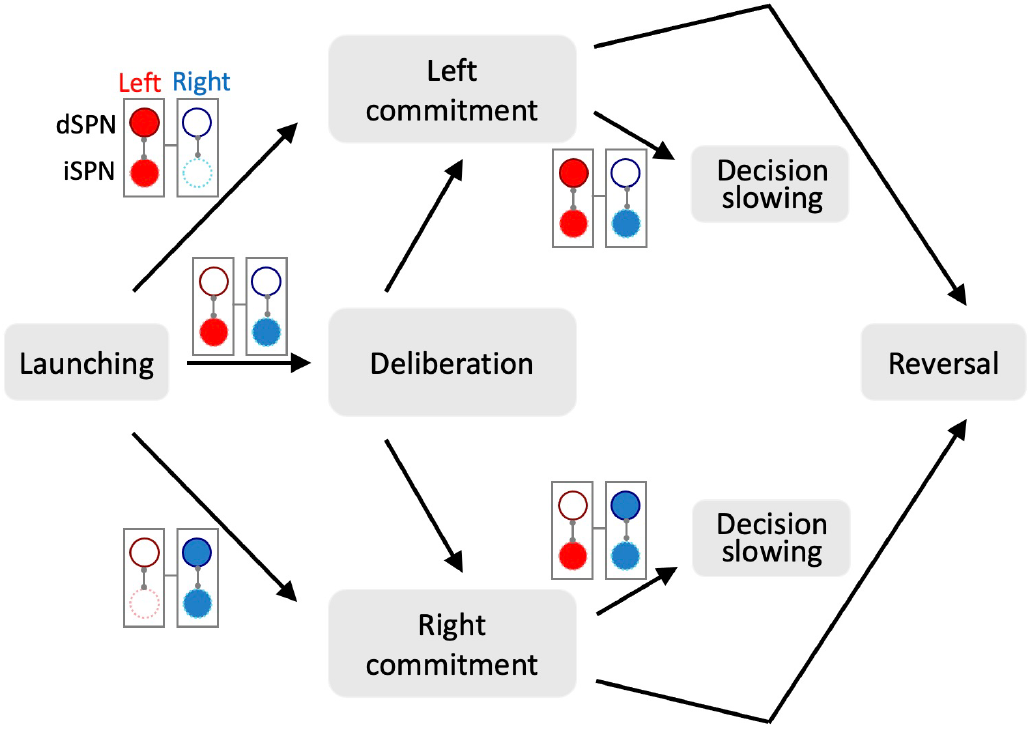
Schematic summary of the three state-dependent roles of iSPNs and their mechanistic interpretation. Red and blue denote left and right action channels, respectively. Upper circles with solid dark outlines represent dSPNs, and lower circles with dotted light outlines represent iSPNs; filled and open circles indicate relatively high and low activity, respectively. The gray blocks summarize decision phases identified in the CLAW analysis (Fig. 3). (1) Preserving uncertainty (launching to deliberation): iSPN activity without dSPN engagement characterizes deliberation and relatively weak choice bias, explained by pliancy engagement that sustains the decision threshold. (2) Signifying commitment (launching to left/right commitment): same-channel dSPN+iSPN coactivation marks increased commitment, driven by the combined influence of choice and responsiveness. (3) Slowing decisions (left/right commitment to decision slowing): subsequent recruitment of opposing-channel iSPNs is associated with longer decision times, explained by reduced choice-ensemble drive that slows evidence accumulation.

Another distinct circuit mechanism emerged during same-channel dSPN+iSPN coactivation. The increased commitment associated with the same-channel iSPN activity was explained by the joint influence of *responsiveness* and *choice*. The choice influence, driven by activity differences between action channels across the direct and indirect pathways, continued to increase toward the selected action along the outer paths (arrow colors for transitions 4→5 and 8→10 in Fig. 5C) and thus strengthened the corresponding rightward or leftward drift drive (*SI Appendix*, Fig. S9B). At the same time, responsiveness, which mainly depends on the global activity of the corticothalamic and direct pathways, became more strongly engaged (Fig. 5B), leading to a further decrease in boundary height (*SI Appendix*, Fig. S9A). Thus, these influences affected different parts of the decision process: choice strengthened drift toward the initially dominant action channel, while responsiveness reduced the amount of evidence needed to reach a decision, which together consolidated the emerging bias. In this context, iSPN activity contributed to commitment within a circuit state where both the direction and the speed of decision formation favored the selected action. This role is illustrated schematically by the routes from launching to left and right commitment in Fig. 6.

Finally, opponent-channel iSPN activation late in the decision process corresponded to small increases in activation of the pliancy and responsiveness ensembles and a decrease in engagement of the choice ensemble. The routes from left and right commitment to decision slowing in Fig. 6 illustrate this SPN activity configuration. The largest impact on decision time was derived from the *choice* ensemble effect (arrow colors for transitions 5→7 and 10→11 in Fig. 5C). The decrease in choice engagement arose because at the coactivation state, the trajectory had already established a dominant action channel, and subsequent activation of the opponent-channel iSPNs reduced the activity difference between the two competing channels. That is, along the left path, increased right-channel iSPN activity reduced the L−R difference; along the right path, increased left-channel iSPN activity reduced the R−L difference. As a result, the choice ensemble’s directional drive regressed back towards 0 (compare arrow colors for transitions 5→7 relative to 4→5, and 10→11 relative to 8→10). Correspondingly, this shift in the choice drive decreased the drift rate (*SI Appendix*, Fig. S9B), slowing evidence accumulation toward the decision boundary and consequently extending decision times.

## Discussion

It is becoming increasingly clear that neural computation arises from low-dimensional interactions among a large number of cell populations, making a one-to-one mapping between anatomically defined cell types and specific functions unrealistic. Cell identity constrains connectivity and physiological capabilities, but it does not uniquely determine the computation expressed at a given moment (43). Rather, computation emerges from the collective state of a neural circuit and from how that state evolves during behavior. This principle is evident in cortical systems, where complex single-neuron responses become interpretable as components of population dynamics that reorganize with behavioral context (44, 45). In the basal ganglia, this perspective calls for moving beyond the canonical description of the direct and indirect pathways as simple Go and No-Go signals (17, 18). Both pathways can make distinct yet coordinated contributions to computation as decisions unfold (21, 23, 24, 27, 28, 30). The central question therefore shifts from what a pathway does in isolation to how evolving activation patterns within each pathway and interactions across pathways jointly shape the decision-making process.

Our study reconciles this dynamical perspective with the canonical pathway organization of the basal ganglia. A labeled CBGT model enabled model-guided inference of putative pathway and action-channel identities in empirical recordings, providing a reference frame for understanding contributions to decision-making dynamics. Across simulations and empirical recordings, iSPN activity was associated with distinct roles at different stages within each decision trajectory: maintaining uncertainty, solidifying commitment, and slowing decisions (Figs. 3 and 4). Projection of activation patterns onto a set of control ensembles further linked these roles to distinct modes of circuit-level regulation of evidence accumulation (Fig. 5), where uncertainty was associated primarily with the pliancy ensemble, commitment with the combined influence of the choice and responsiveness ensembles, and decision slowing with the choice ensemble. These findings highlight that pathway identity and circuit structure constrain the range of possible circuit interactions, whereas evolving population dynamics determine how those interactions are expressed during each decision.

This view refines rather than rejects the canonical direct/indirect pathway account. In the striatal CLAW (Fig. 3), dSPN activity provided the clearest choice-specific structure (outer CLAW), while the iSPN-dominated trajectories were associated with uncertainty before choice-related asymmetry emerged (inner CLAW). Once an action channel became dominant, however, same-channel iSPN recruitment accompanied commitment, whereas opponent-channel iSPN recruitment weakened the existing asymmetry and slowed the decision. Thus, the two path-ways retain functional distinctions, but their behavioral impacts depend on when they are recruited along an evolving decision trajectory.

Although the empirical CLAW preserved the principal topology predicted by the CBGT model, it was not perfectly symmetric between the two action populations. Compared with the CBGT CLAW, the empirical CLAW exhibited three directional asymmetries: (i) trajectories launched more frequently into left-channel states (*P* (0 → 8) = 0.21 versus *P* (0 → 4) = 0.15); (ii) the bilateral-iSPN state reached along the inner deliberative route showed a bias toward right choices (state 3, left-choice probability = 0.38); and (iii) outer states associated with the right channel had longer decision times than the corresponding left-channel states (state 4: 250 ms versus state 8: 232 ms; state 5: 250 ms versus state 10: 228 ms). These asymmetries may reflect influences that were absent from the CBGT model but present in the empirical task, behavior, and recording design. In the IBL task, the prior probability of stimulus location alternated between uncued blocks. Mice thus formed a subjective estimate of this prior primarily from their recent actions; the influence of prior information on choice was especially strong when visual evidence was weak or absent (46). Mice in the IBL task also alternated among engaged and biased behavioral states that persisted for tens to hundreds of trials (47). Such latent biases can affect both the probability and timing of a decision: prior information can produce more frequent and faster choices toward a favored alternative, largely through a shift in the starting point of evidence accumulation (48), whereas recent choice history can modify the rate of subsequent evidence accumulation (49). These effects could change which paths are entered, how long deliberation persists, and how quickly each choice is reached.

Hence, one possible interpretation within the empirical CLAW is that trials ending in left choices more often entered the left outer path directly from the launching state, whereas trials ending in right choices more often remained within the inner deliberative route before reaching the right outer path. This unequal routing could account for more frequent launching into state 8, the relative enrichment of subsequent right choices among trials reaching state 3, and the longer decision times for trials that visited the outer right states. Importantly, our CLAW analysis on empirical recordings intentionally matched left and right choice trials within stimulus contrast bins, yielding equal numbers of final left and right choices. The observed asymmetries therefore cannot be attributed to unequal choice counts, but instead reflect differences in the paths and timing through which the two choices were reached. In line with this interpretation, stochastic competition between neural populations can generate idiosyncratic directional tendencies even under symmetric experimental conditions (50), providing one possible source of the unequal routing observed here. In addition, the IBL recordings sampled the left forebrain and midbrain, including the striatal regions analyzed in this work, which typically represent contralateral stimuli and actions (32). This hemispheric sampling may have further contributed to unequal empirical representations of the two action channels, although it does not by itself predict the direction of the observed differences. Thus, the empirical asymmetries do not contradict the common CLAW topology and associated decision outcomes, but illustrate how prior information, latent behavioral states, intrinsic neural variability, and recording laterality may affect their expression in biological data.

These considerations also motivate direct experimental tests of the state-context framework. A key limitation of the present study is that pathway identities in the empirical recordings were inferred from extracellular activity rather than identified molecularly. Recent studies have directly determined neuron identities during behavior, including probabilistic action selection, sustained action suppression, and hesitation under uncertainty, using optogenetic and electrophysiological identification or photoidentified recordings (30, 51, 52). These studies establish that pathway membership can be identified and manipulated during decision-related behavior. Applying these methods to the visual two-choice task, ideally with bilateral recordings, would test whether identified dSPNs and iSPNs reproduce the inferred activity patterns and separate into the functionally defined left and right action populations predicted by the model.

Such experiments would also enable tests of the three proposed roles of iSPN activity through observations and perturbations in the corresponding circuit states. First, perturbing iSPNs during iSPN-alone states could test whether their activity maintains an undecided state and delays commitment. Second, if selected-channel iSPN recruitment signifies growing commitment, recordings should show iSPN activation rising until decision time as evidence increasingly favors that option, with dSPNs in the same channel already above the activation threshold. Weakening the evidence for that option sufficiently to keep iSPN activation in that channel subthreshold should increase the probability of a choice reversal or a delayed response. Finally, suppressing iSPNs recruited in the opposing channel should shorten decision times if those neurons contribute to decision slowing. Because these predictions depend on the circuit state in which iSPN activity occurs, triggering perturbations when specific patterns of population activity emerge would test them more directly than aligning them only to a cue or movement onset.

Nonetheless, our findings provide direct support for the idea that engagement of a specific cell population does not map to a singular function, but depends on patterns of coactivation across the entire network. Synaptic pathways define the interactions available to a circuit, while the evolving pattern of collective activity determines which interactions are behaviorally relevant at a given moment. This perspective complements emerging views of neural computation, in which roles are flexible and collectively defined, and may extend beyond the basal ganglia. Wherever anatomically defined populations interact to shape behavior, their functional roles may shift over time and across behavioral contexts.

## Materials and Methods

We summarize the analysis design and principal procedures below; full methodological and implementation details are provided in the *SI Appendix*.

### Data Sources and Analysis Sequence

Simulated CBGT activity came from 300 independently parameterized networks (34) in a conductance-based spiking CBGT model implemented in CBGTPy (33). Each network contributed 50 trials and four labeled SPN populations. Empirical analyses used caudoputamen/dorsal-striatal (CP) Neuropixels recordings from the Steinmetz and International Brain Laboratory (IBL) visual two-choice datasets (31, 32). Spikes were aligned to movement onset, binned at 10 ms, causally smoothed, and normalized to pre-stimulus firing. After applying the initial inclusion criteria at the trial, unit, and session levels, 11 recording sessions were retained for clustering.

The analysis proceeded in the following sequence. We first used the CBGT model, where SPN channel and pathway identities were known, to identify activity features that distinguished these groups. Guided by those features, we clustered empirical units to infer their action channel and pathway labels. Shuffle controls and the assessment of potential FSI inclusion were then conducted among the clustered units. Finally, we excluded units that met the putative FSI criteria and used the remaining units, with their inferred SPN subtype labels, for CLAW construction and prediction analyses.

### Two-Stage SPN Subtype Inference

Stage 1 inferred action channel identity from mean pre-decision activity. This stage used z-scored 12-dimensional vectors formed from the final six pre-decision time bins averaged separately over left-choice and right-choice trials. Ward-linkage agglomerative clustering (53) divided units into two action channels. For the simulated data, cluster identities were aligned with the known labels using the Hungarian algorithm (54). A forward bin-inclusion analysis evaluated how the number of pre-decision bins affected recovery of the known CBGT action channel labels, supporting the use of six bins per choice in Stage 1 (*SI Appendix*, Fig. S2).

Stage 2 inferred pathway identity within each Stage 1 action channel using trial-level correlation structure. In the simulated data, firing rate profiles of the dSPN population were observed to have a weaker mean correlation with those of the two opposite-channel populations than the profiles of the iSPN population. In the empirical data, pairwise Pearson correlations, *r*, between neuronal firing rate vectors were converted to correlation distances, *D* = 1 − *r*. Agglomerative hierarchical clustering with average linkage (55) was then applied separately within each inferred action channel, dividing the units into two subclusters. To assign pathway identities to subclusters, trials within each session were ranked by decision time and split into fast and slow halves. Among the four possible cross-channel subcluster pairings, the pair showing the largest decrease in correlation from fast to slow trials was labeled as the putative dSPN pair; the remaining pair was labeled as the putative iSPN pair.

### Assessment of Clustered Empirical Units

Three shuffle controls independently permuted within-unit interspike intervals, choice labels, or fast/slow trial labels. We evaluated the original empirical data and each corresponding shuffled replicate against prespecified criteria derived from the simulated model’s predictions. For each session, we calculated the percentage of criteria satisfied by the original data and by each shuffled replicate.

Potential FSI inclusion was assessed after the empirical subtype labels had been assigned. Waveform width at half of the maximum negative deflection was measured from the peak-amplitude channel and paired with whole-session firing rate. Units with a waveform width *<* 0.15 ms and a whole-session firing rate *>* 10 Hz were flagged as putative FSIs (37–39). These flagged units were removed; all the other units and their subtype labels were retained for the subsequent CLAW and prediction analyses.

### Striatal CLAW Construction and Prediction Tests

Based on our previously established Circuit Logic Assessed via Walks (CLAW) framework (34, 40), we represented continuous striatal activity traces as trajectories through a discrete state space. At each 10-ms time bin, activity was encoded as a binary vector *s*_*k*_ = [dL, dR, iL, iR], where each element indicated whether the corresponding SPN group (d for direct, i for indirect, L for left channel, R for right channel) was active or not. Each element was assigned a value of 1 when the corresponding population firing rate exceeded its binarization threshold and 0 otherwise (see thresholds in *SI Appendix*, Fig. S1D). Consecutive time bins assigned to the same state were collapsed; adjacent states in the compressed sequence were counted as transitions; and the final state prior to decision was followed by an END event. Transition probability was calculated over all outgoing events, including END; terminal probability was *P* (END | state). We calculated left choice probability (node color) and mean decision time (gray bar) across trials that visited each state. Low-frequency edges were omitted only from the visualization.

Predictions A and B in Fig. 4 used 5,000 trial-level bootstrap resamples, with group differences assessed using two-sided independent-samples *t*-tests; prediction A additionally used a one-way ANOVA and Bonferroni correction. Prediction C used two-sided independent-samples *t*-tests on the raw decision time distributions. Model-derived and empirical data were analyzed separately.

### Control Ensemble and DDM Projection

The DDM parameters and the CBGT activity for the 300 network configurations were used to identify control ensembles through canonical correlation analysis (CCA), as in prior work (27, 34, 41). For each network, distributions of choice and decision time were fit with a static drift-diffusion model, yielding the corresponding configuration of static DDM parameters. CCA was then applied to relate these parameters to an 18-dimensional representation of CBGT activity, consisting of the firing rates of two shared neuron populations, 8 across-channel summed firing rates, and 8 between-channel differences in firing rates. The resulting canonical variates defined correlated patterns of CBGT activity and DDM parameters, and the three leading components defined the pliancy, responsiveness, and choice ensembles analyzed here (reduced circuit, Fig. 5A, left subpanels; full circuit, *SI Appendix*, Fig. S8).

For each network and CLAW state, activity across the full set of CBGT populations was averaged over all pre-decision bins assigned to that state. For each transition, the change in circuit activity from the source to the target state was projected onto the neural loadings of the choice, responsiveness, and pliancy ensembles and then onto their associated DDM parameters. Transition effects were summarized across networks in which both states occurred. In (34), we evaluated this use of static CCA axes with toy-model generalization tests, which supported a shared mapping at the time-bin and zone scales when the neural activity-policy relationship remained stable. This analysis was restricted to the model because the empirical recordings did not provide simultaneous activity traces and identification of cell types for the full set of CBGT populations.

## Data, Materials, and Software Availability

The codebase for the CBGT computational model is publicly available at https://github.com/CoAxLab/CBGTPy. The original simulated CBGT datasets and corresponding DDM parameter configurations used in this study are available at https://github.com/zhuojunyu-appliedmath/CLAW. Analysis code and derived data supporting this study, including analyses of publicly available experimental datasets from the Steinmetz and International Brain Laboratory groups, are available at https://github.com/zhuojunyu-appliedmath/context-dependent-role-iSPNs.

## ACKNOWLEDGMENTS

We thank Eric Yttri, Cati Vich Llompart, Jyotika Bahuguna, and all members of the exploratory intelligence group for their helpful comments on this work. This work was supported by NIH award R01DA059993 as part of the CRCNS program. JER is also partly supported by NIH award R01NS125814, also part of the CRCNS program.

## References

1. T Willis, Cerebri anatome: Cui accessit nervorum descriptio et usus. (Typis Ja. Flesher, impensis Jo. Martyn & Ja. Allestry, London), (1664).

2. GE Alexander, MR DeLong, PL Strick, Parallel organization of functionally segregated circuits linking basal ganglia and cortex. Annu. review neuroscience 9, 357–381 (1986).

3. GE Alexander, MD Crutcher, Functional architecture of basal ganglia circuits: neural substrates of parallel processing. Trends neurosciences 13, 266–271 (1990).

4. JW Mink, The basal ganglia: focused selection and inhibition of competing motor programs. Prog. neurobiology 50, 381–425 (1996).

5. K Gurney, TJ Prescott, P Redgrave, A computational model of action selection in the basal ganglia. i. a new functional anatomy. Biol. cybernetics 84, 401–410 (2001).

6. M Desmurget, RS Turner, Motor sequences and the basal ganglia: kinematics, not habits. The J. Neurosci. 30, 7685–7690 (2010).

7. RM Costa, D Cohen, MA Nicolelis, Differential corticostriatal plasticity during fast and slow motor skill learning in mice. Curr. Biol. 14, 1124–1134 (2004).

8. HH Yin, et al., Dynamic reorganization of striatal circuits during the acquisition and consolidation of a skill. Nat. neuroscience 12, 333–341 (2009).

9. MT Ullman, Contributions of memory circuits to language: The declarative/procedural model. Cognition 92, 231–270 (2004).

10. M Teichmann, E Dupoux, S Kouider, AC Bachoud-Lévi, The role of the striatum in processing language rules: evidence from word perception in huntington’s disease. J. cognitive neuroscience 18, 1555–1569 (2006).

11. VS Chakravarthy, D Joseph, RS Bapi, What do the basal ganglia do? a modeling perspective. Biol. cybernetics 103, 237–253 (2010).

12. R Bogacz, K Gurney, The basal ganglia and cortex implement optimal decision making between alternative actions. Neural computation 19, 442–477 (2007).

13. Y Niv, ND Daw, D Joel, P Dayan, Tonic dopamine: opportunity costs and the control of response vigor. Psychopharmacology 191, 507–520 (2007).

14. EA Yttri, JT Dudman, Opponent and bidirectional control of movement velocity in the basal ganglia. Nature 533, 402–406 (2016).

15. GS Berns, TJ Sejnowski, A computational model of how the basal ganglia produce sequences. J. cognitive neuroscience 10, 108–121 (1998).

16. O Hikosaka, et al., Parallel neural networks for learning sequential procedures. Trends neurosciences 22, 464–471 (1999).

17. RL Albin, AB Young, JB Penney, The functional anatomy of basal ganglia disorders. Trends neurosciences 12, 366–375 (1989).

18. MR DeLong, Primate models of movement disorders of basal ganglia origin. Trends neurosciences 13, 281–285 (1990).

19. N Mallet, et al., Dichotomous organization of the external globus pallidus. Neuron 74, 1075–1086 (2012).

20. M Cazorla, et al., Dopamine d2 receptors regulate the anatomical and functional balance of basal ganglia circuitry. Neuron 81, 153–164 (2014).

21. J Bahuguna, A Aertsen, A Kumar, Existence and control of go/no-go decision transition threshold in the striatum. PLoS computational biology 11, e1004233 (2015).

22. S Bariselli, W Fobbs, M Creed, A Kravitz, A competitive model for striatal action selection. Brain research 1713, 70–79 (2019).

23. K Dunovan, T Verstynen, Believer-skeptic meets actor-critic: rethinking the role of basal ganglia pathways during decision-making and reinforcement learning. Front. neuroscience 10, 106 (2016).

24. F Tecuapetla, X Jin, SQ Lima, RM Costa, Complementary contributions of striatal projection pathways to action initiation and execution. Cell 166, 703–715 (2016).

25. W Wei, JE Rubin, XJ Wang, Role of the indirect pathway of the basal ganglia in perceptual decision making. J. Neurosci. 35, 4052–4064 (2015).

26. H Li, X Jin, Multiple dynamic interactions from basal ganglia direct and indirect pathways mediate action selection. Elife 12, RP87644 (2023).

27. C Vich, M Clapp, JE Rubin, T Verstynen, Identifying control ensembles for information processing within the cortico-basal ganglia-thalamic circuit. PLOS Comput. Biol. 18, e1010255 (2022).

28. G Cui, et al., Concurrent activation of striatal direct and indirect pathways during action initiation. Nature 494, 238–242 (2013).

29. F Tecuapetla, S Matias, GP Dugue, ZF Mainen, RM Costa, Balanced activity in basal ganglia projection pathways is critical for contraversive movements. Nat. communications 5, 4315 (2014).

30. S Nonomura, et al., Monitoring and updating of action selection for goal-directed behavior through the striatal direct and indirect pathways. Neuron 99, 1302–1314 (2018).

31. NA Steinmetz, P Zatka-Haas, M Carandini, KD Harris, Distributed coding of choice, action and engagement across the mouse brain. Nature 576, 266–273 (2019).

32. International Brain Laboratory, et al., A brain-wide map of neural activity during complex behaviour. Nature 645, 177–191 (2025).

33. M Clapp, et al., Cbgtpy: An extensible cortico-basal ganglia-thalamic framework for modeling biological decision making. PLoS One 20, e0310367 (2025).

34. Z Yu, T Verstynen, JE Rubin, How the dynamic interplay of cortico-basal ganglia-thalamic pathways shapes the time course of deliberation and commitment. PLOS Comput. Biol. 22, e1012966 (2026).

35. S Amadio, et al., Motor evoked potentials in a mouse model of chronic multiple sclerosis. Muscle & Nerve: Off. J. Am. Assoc. Electrodiagn. Medicine 33, 265–273 (2006).

36. V Castoldi, E Rossi, S Marenna, G Comi, L Leocani, Improving reproducibility of motor evoked potentials in mice. J. Neurosci. Methods 367, 109444 (2022).

37. JM Burkhardt, X Jin, RM Costa, Dissociable effects of dopamine on neuronal firing rate and synchrony in the dorsal striatum. Front. integrative neuroscience 3, 562 (2009).

38. JD Berke, Uncoordinated firing rate changes of striatal fast-spiking interneurons during behavioral task performance. The J. Neurosci. 28, 10075–10080 (2008).

39. GJ Gage, CR Stoetzner, AB Wiltschko, JD Berke, Selective activation of striatal fast-spiking interneurons during choice execution. Neuron 67, 466–479 (2010).

40. Z Yu, JE Rubin, T Verstynen, Learning reweights the decision dynamics of cortico-basal ganglia-thalamic pathways from deliberation to commitment. bioRxiv (2026).

41. J Bahuguna, T Verstynen, JE Rubin, How cortico-basal ganglia-thalamic subnetworks can shift decision policies to increase reward rate. PLOS Comput. Biol. 21, e1013712 (2025).

42. R Ratcliff, A theory of memory retrieval. Psychol. review 85, 59 (1978).

43. DC Burnston, TJ Ryan, An alternative to encoding for thinking about neural representation. Philos. Mind Sci. 7 (2026).

44. MM Churchland, et al., Neural population dynamics during reaching. Nature 487, 51–56 (2012).

45. V Mante, D Sussillo, KV Shenoy, WT Newsome, Context-dependent computation by recurrent dynamics in prefrontal cortex. nature 503, 78–84 (2013).

46. C Findling, et al., Brain-wide representations of prior information in mouse decision-making. Nature 645, 192–200 (2025).

47. ZC Ashwood, et al., Mice alternate between discrete strategies during perceptual decision-making. Nat. Neurosci. 25, 201–212 (2022).

48. MJ Mulder, EJ Wagenmakers, R Ratcliff, W Boekel, BU Forstmann, Bias in the brain: a diffusion model analysis of prior probability and potential payoff. J. Neurosci. 32, 2335–2343 (2012).

49. AE Urai, JW De Gee, K Tsetsos, TH Donner, Choice history biases subsequent evidence accumulation. elife 8, e46331 (2019).

50. L Lebovich, R Darshan, Y Lavi, D Hansel, Y Loewenstein, Idiosyncratic choice bias naturally emerges from intrinsic stochasticity in neuronal dynamics. Nat. human behaviour 3, 1190–1202 (2019).

51. BF Cruz, et al., Action suppression reveals opponent parallel control via striatal circuits. Nature 607, 521–526 (2022).

52. MA Geramita, SE Ahmari, EA Yttri, The striatal indirect pathway mediates hesitation. Nat. Neurosci. 29, 287–292 (2026).

53. JH Ward Jr, Hierarchical grouping to optimize an objective function. J. Am. statistical association 58, 236–244 (1963).

54. HW Kuhn, The hungarian method for the assignment problem. Nav. research logistics quarterly 2, 83–97 (1955).

55. GN Lance, WT Williams, A general theory of classificatory sorting strategies: 1. hierarchical systems. Comput. J. 9, 373–380 (1967).

