## Supporting Information Appendix for "How striatal neurons influence decision-making depends on broader circuit context: evidence from theory-driven cell type identification"

**This PDF file includes:**

Supporting text
Figs. S1 to S9
SI References

### Supporting Information Text

### CBGT Model Simulations

Simulated activity was obtained from a previously established biologically grounded spiking model of the CBGT circuit, implemented in CBGTPy (1). The model comprised 10 neuronal populations: cortical interneurons (CxI), excitatory cortical neurons (Cx), direct- and indirect-pathway spiny projection neurons (dSPNs and iSPNs), striatal fast-spiking interneurons (FSIs), prototypical and arkypallidal neurons of the external globus pallidus (GPeP and GPeA), the subthalamic nucleus (STN), the internal globus pallidus (GPi), and the thalamus (Th) (Fig. S1A). CxI and FSI populations were shared across channels; the other populations were represented separately in left and right channels.

Each simulated trial was initiated by a sustained stimulus input to the cortical neurons in both channels, producing stochastic cortical activity that propagated through the striatum and downstream basal ganglia nuclei. A decision occurred when the instantaneous firing rate of either left- or right-channel thalamic population first reached 30 Hz (Fig. S1B). Individual neurons followed conductance-based integrate-and-fire-or-burst dynamics (see model equations in (1)).

The simulated dataset, originally generated in (2), comprised 300 independently parameterized CBGT networks produced using a genetic algorithm to sample distinct configurations of synaptic connection strengths (see also (3)). Each network was simulated for 50 trials. Networks were divided into equal-count fast, intermediate, and slow tertiles according to median decision time (Fig. S1C). Because both pathway identity and action-channel assignment were known for every modeled SPN population, the 300 networks yielded 1,200 labeled model SPN units, providing a ground-truth reference for developing and evaluating the subtype-inference procedure in this study. A complete description of the network architecture, model equations, neuronal dynamics, and simulation procedures is provided in (1, 2).

### Empirical Datasets and Preprocessing

We analyzed two public Neuropixels recordings from mice performing visual two-choice tasks (4, 5). Units assigned to caudoputamen/dorsal striatum (CP) and the corresponding eligible behavioral trials were retained according to the source datasets. No new animal experiments were conducted.

Spikes were aligned to detected movement onset and counted in 10-ms bins. Firing rates were smoothed using a causal half-Gaussian kernel ($\sigma = 25$ ms) and divided by each unit's mean firing rate during the 200 ms preceding stimulus onset plus 1 spike/s. Here we approximated the decision time as 10 ms before detected movement onset. This offset placed the analysis window before measurable motor execution and was consistent with reported cortical motor-evoked potential latencies in mice (6, 7).

Unit quality control followed the criteria of the source datasets. We additionally excluded units if their 12-dimensional pre-decision firing rate pattern was nearly flat (defined as a maximum-to-minimum difference below 0.05) or if the fitted slope was negative for either choice condition. A session entered clustering only if it contained at least eight eligible striatal units. After filtering, 11 sessions (4 from the Steinmetz dataset and 7 from the IBL dataset) were retained for analysis.

### SPN Subtype Inference

The inference approach was developed using the simulated CBGT data, for which ground-truth labels were known, and subsequently applied to the empirical recordings. Stage 1 inferred action-channel identity from mean activity before decision. For each unit, firing rate activity in the final six 10-ms bins before the decision cutoff was averaged separately across left- and right-choice trials and concatenated into a 12-dimensional vector. Each vector was z-scored across dimensions, and agglomerative hierarchical clustering with Euclidean distance and Ward linkage (8) partitioned the units into two clusters. In simulated data, arbitrary cluster numbers were aligned to the known action channels with the Hungarian algorithm (9) to find the one-to-one mapping that maximized agreement between cluster assignments and ground-truth labels. ARI itself is invariant to this alignment. In empirical data, where ground-truth channel identities were unavailable, the cluster with the larger mean left-choice minus right-choice activity difference was assigned to the putative left channel, and the other cluster to the putative right channel.

To evaluate the six-bin choice of feature window, Stage 1 was repeated on the 1,200 modeled SPN populations using forward inclusion from the earliest bin. The first analysis used bin $-6$ from each choice condition, producing two dimensions; subsequent analyses added bins $-5$ through $-1$ sequentially, producing 4, 6, 8, 10, and 12 dimensions. Every subset was selected from the same z-scored complete activity matrix, and the same two-cluster Ward procedure was applied. ARI was calculated against the known left/right channel identities. The complete 12-dimensional feature achieved ARI $= 0.983$ and exceeded every shorter forward subset (Fig. S2).

Stage 2 distinguished pathway identity within each Stage 1 action channel using trial-level correlation structure. In the model, firing rates from the final six pre-decision bins were concatenated across trials and z-scored within network. Pairwise Pearson correlations were then computed among the four SPN populations using the same trials in the same order. Within each inferred action channel, the population with weaker mean correlation to the two opposite-channel populations was labeled dSPN; the other was labeled iSPN. The complete four-subtype assignment achieved ARI $= 0.873$ against the known model labels (Fig. 1D in the main text).

In empirical sessions, correlations were estimated over a wider peri-movement interval ($-100$ to $+50$ ms) to obtain more stable correlation estimates from noisy extracellular recordings. Pairwise Pearson correlations, $r$, between the resulting firing

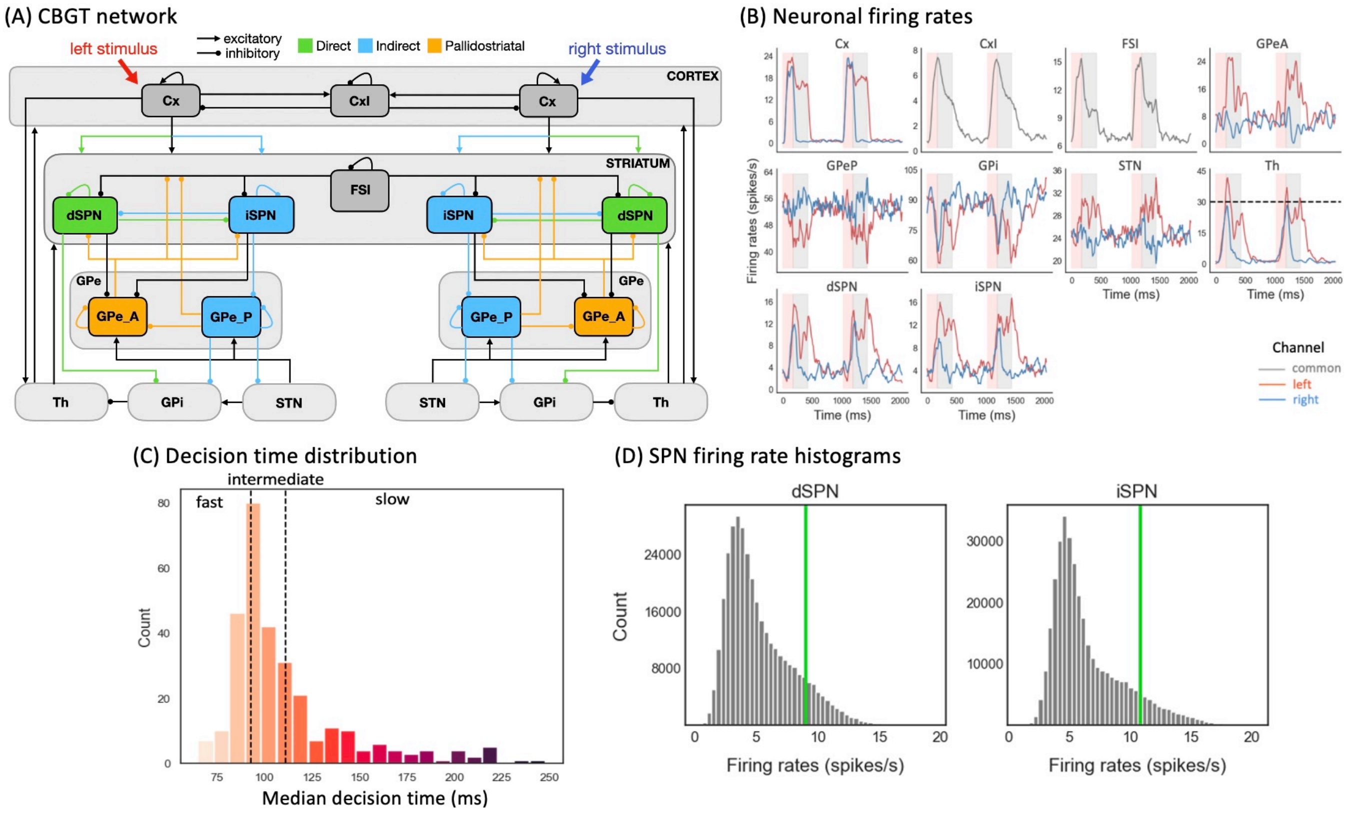


**Fig. S1.** (A) Schematic of the two-channel cortico-basal ganglia-thalamic (CBGT) circuit used to model competition between left- and right-stimulus alternatives. Colored populations and projections indicate the direct pathway (green), indirect pathway (blue), and pallidostriatal pathway (gold). Note that the colors highlight the key nuclei shaping each pathway's influence, rather than every anatomical component. For example, internal globus pallidus (GPi) participates in both direct and indirect pathways, and subthalamic nucleus (STN) is part of the indirect pathway, but we color only the nuclei whose activity most clearly reflects each pathway's role during decisions. Arrows ending in dots indicate postsynaptic sites of inhibitory connections, while arrows ending in triangles indicate excitatory connections. Cx, cortical neurons; CxI, inhibitory cortical interneurons; FSI, fast spiking interneurons; dSPN, direct spiny projection neurons; iSPN, indirect spiny projection neurons; GPe, external globus pallidus; GPeP, prototypical neurons; GPeA, arkypallidal neurons; GPi, internal globus pallidus; STN, subthalamic nucleus; Th, thalamus. (B) Example population firing rates during two consecutive trials. Red and blue traces correspond to the left and right channels, respectively, and gray traces denote populations shared across channels. Pink regions represent the decision-making phase, occurring before the thalamus of one of the action channels reaches the decision threshold of 30 Hz (dashed black line in Th panel). Grey regions represent the consolidation phase, where partial cortical input to the selected channel is sustained. The unshaded regions represent the inter-trial interval. (C) Distribution of median decision times across the 300 simulated CBGT networks, categorized into three equal-count tertiles defining fast (left), intermediate (middle), and slow (right) networks, demarcated by vertical dashed black lines. (D) Distributions of dSPN and iSPN firing rates across all simulated trials up to decision times. In each panel, the vertical green line indicates the firing rate threshold used for binarization, defined as the 90th percentile of the corresponding firing rate distribution. Figure reproduced from (2).

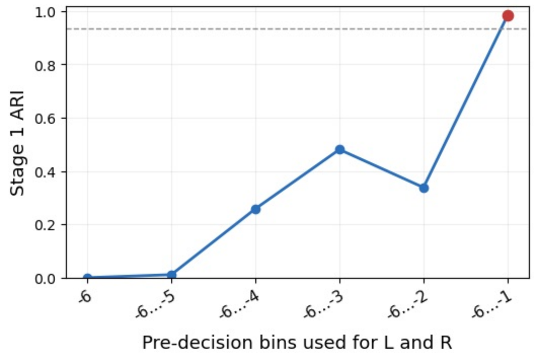


**Fig. S2.** Stage 1 action-channel recovery using different numbers of pre-decision bins. Stage 1 clustering was repeated for 1,200 simulated SPN populations from 300 CBGT networks using progressively longer subsets of the six 10-ms bins preceding the decision for both left-choice and right-choice trials. Starting with bin -6, bins were added sequentially toward the decision at bin -1, increasing the feature dimension from 2 to 12. The adjusted Rand index (ARI) quantifies agreement between the inferred and true action channels. The red point denotes the complete 12-dimensional feature used in the main analysis, which achieved an ARI of 0.983. The horizontal dashed line marks 95% of this value. Channel recovery was substantially more accurate with the complete six-bin representation than with any partial subset.

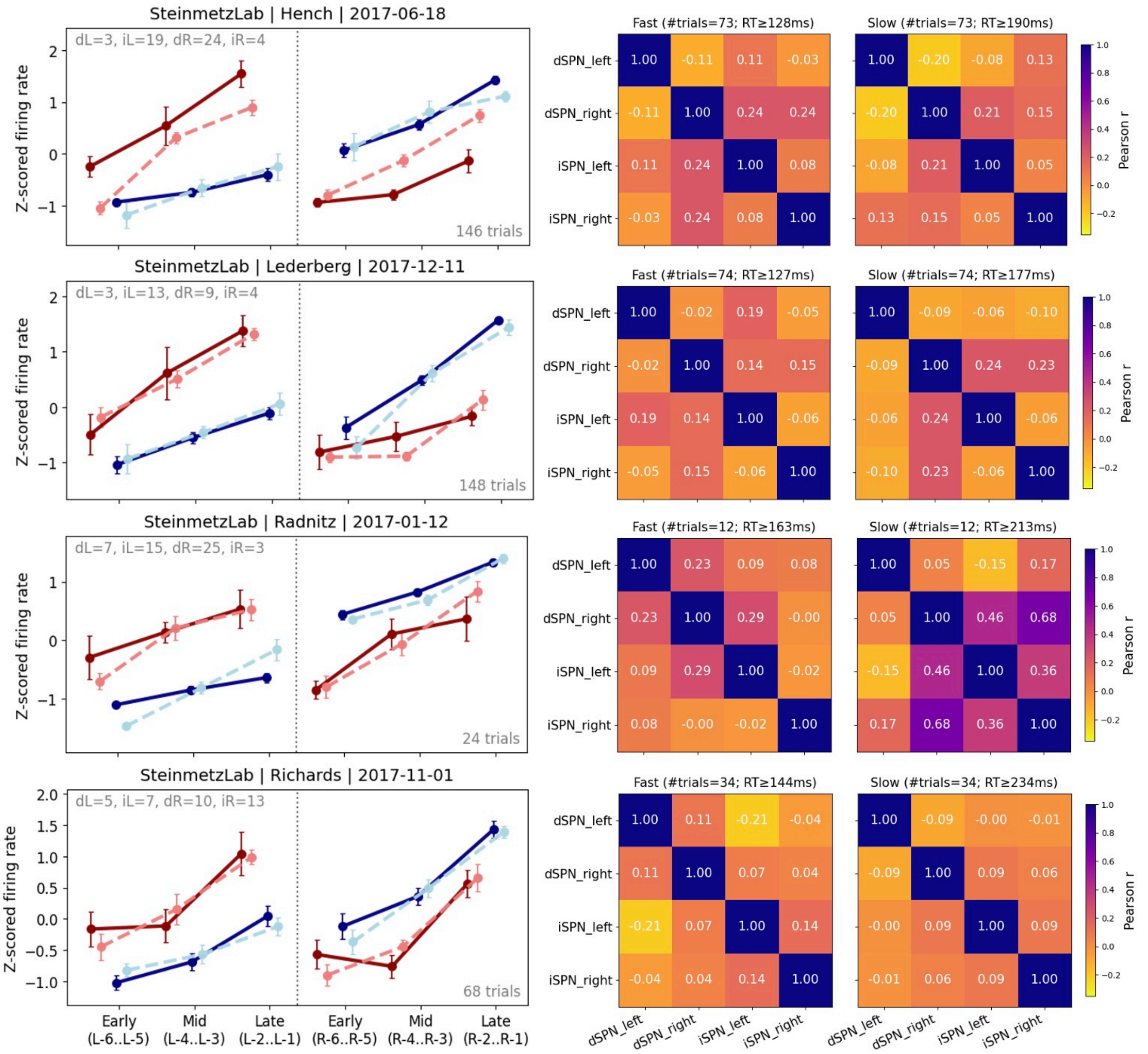


**Fig. S3.** Firing rate profiles (left) and correlation structure (right) of putative SPNs across Steinmetz Laboratory recordings. Putative SPN subtypes were inferred from recordings reported in (4). Each row shows results from one recording session. In the left panels, firing rates were z-scored and averaged across three pre-movement windows: early, middle, and late, corresponding to bins 6–5, 4–3, and 2–1 before the decision, respectively. Color denotes the inferred action channel, and line style denotes the inferred pathway identity. Gray text indicates the number of units assigned to each subtype, and the total number of trials is shown in the lower right of each panel. Error bars indicate SEM. The right panels show Pearson correlation matrices for the firing rates of the four inferred SPN clusters, calculated separately for fast and slow decision trials. The number of trials and decision times for each subset are indicated above each matrix.

rate series were converted to correlation distances, $D = 1 - r$. Average-linkage agglomerative clustering (10) was applied separately within each inferred action channel and cut into two subclusters per channel. To assign the subtype labels, empirical trials were first divided into equal-count fast and slow halves according to their decision times. Among the four cross-channel subcluster pairings, the pair with the largest decrease in correlation from fast to slow trials was labeled the putative dSPN pair; the remaining pair was labeled the putative iSPN pair. The resulting pre-decision firing rate patterns and fast- versus slow-trial correlation matrices are shown for Steinmetz sessions in Fig. S3 for and for IBL sessions in Fig. S4. These action-channel and pathway labels were then fixed for all subsequent analyses.

### Shuffle Controls

Three null controls were performed separately for each of the four Steinmetz sessions, with 50 repetitions per control. The *interspike-interval (ISI) control* independently permuted each unit's ISIs and reconstructed a surrogate spike train, preserving spike count and the unit's marginal ISI distribution while disrupting alignment to task events. The full clustering and subtype-assignment workflow was then repeated. The *choice-label control* permuted left and right labels, preserving the trial

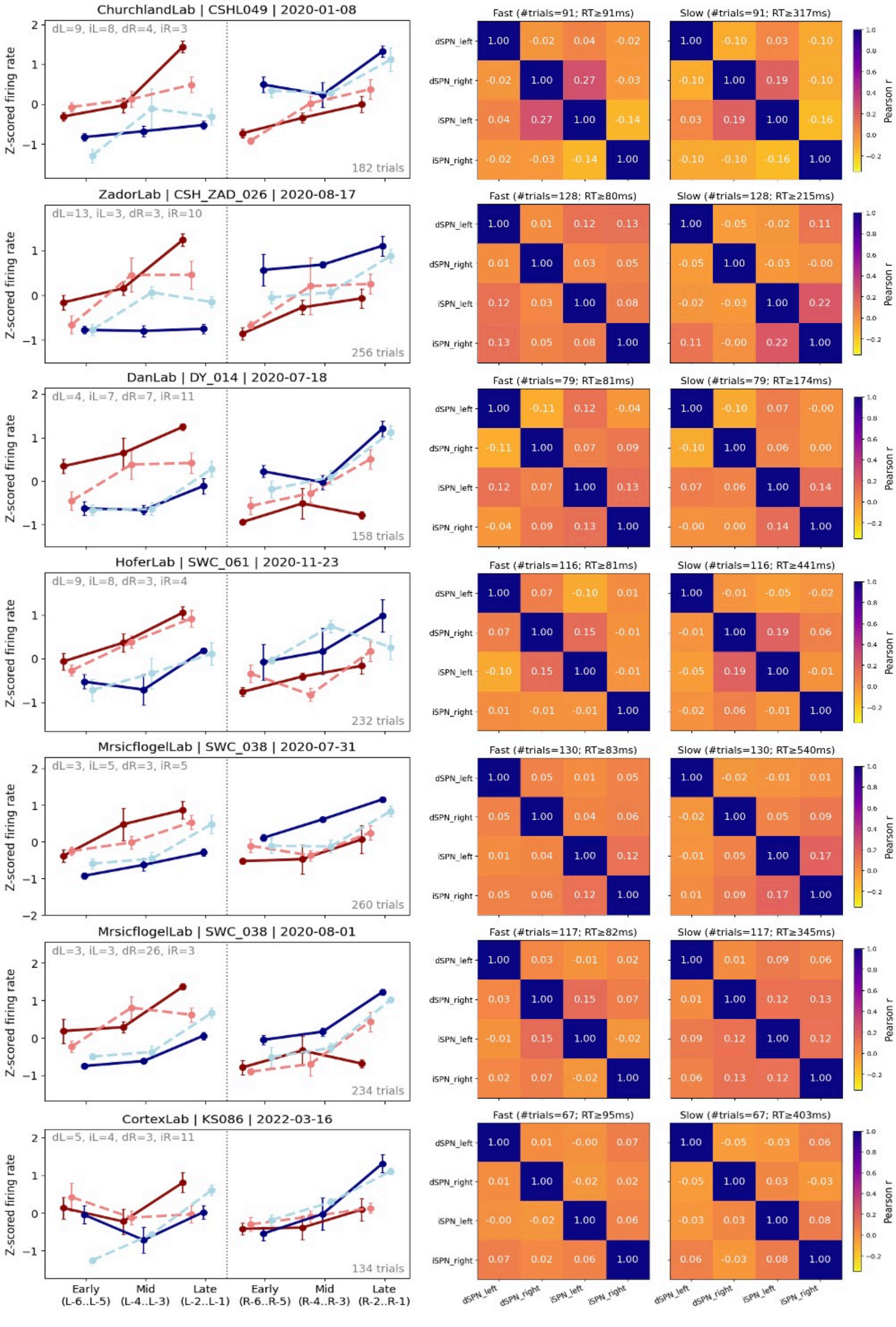

**Fig. S4.** Firing rate profiles (left) and correlation structure (right) of putative SPNs across IBL recordings. Putative SPN subtypes were inferred from recordings reported in (5). Each row shows results from one recording session. Conventions and notation are as in Fig. S3.

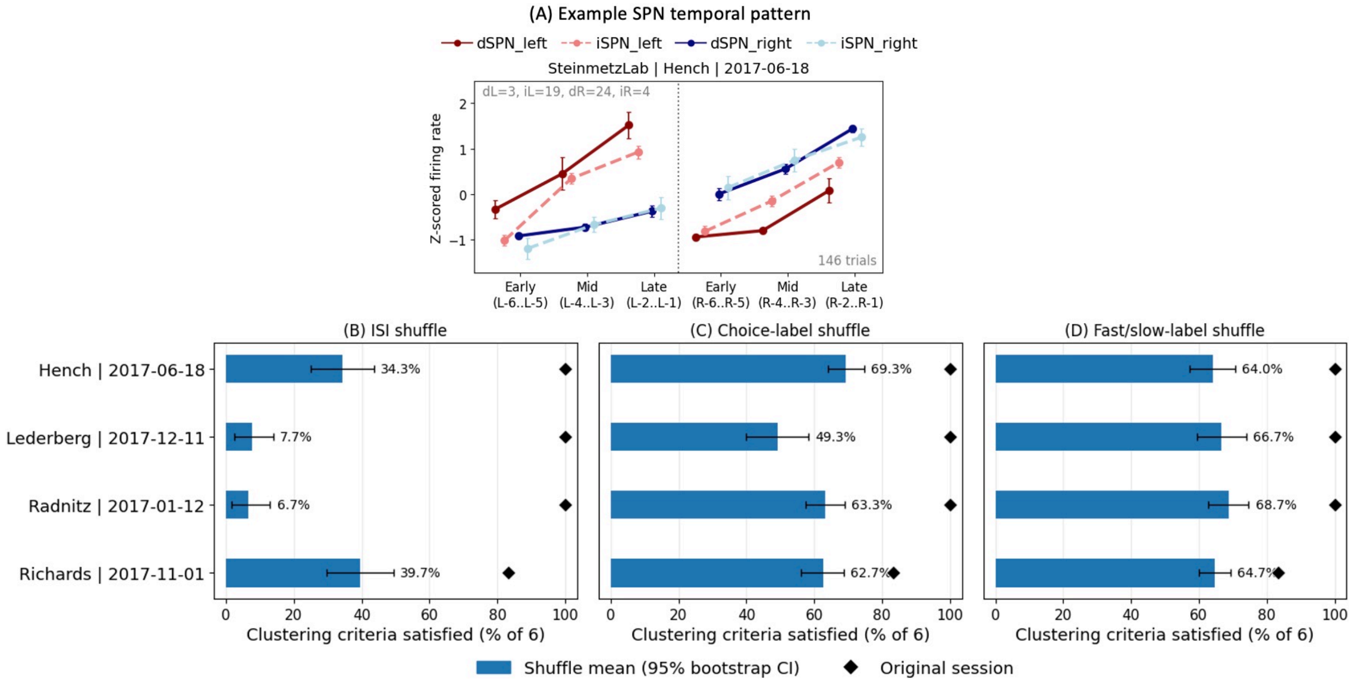


**Fig. S5.** Shuffle control for the empirical SPN subtype classification. (A) Example z-scored SPN firing rate profiles across the early, middle, and late windows preceding left and right choices. (B–D) Percentage of six model-predicted criteria satisfied in each Steinmetz session under three shuffle controls. Blue bars show the mean across 50 shuffle repetitions, with 95% bootstrap confidence intervals; black diamonds show the corresponding values for the original data. (B) Interspike intervals (ISIs) were permuted independently within each unit, preserving spike count and the ISI distribution. (C) Left- and right-choice labels were shuffled within evidence strata. (D) Fast and slow decision labels were shuffled within each choice condition. Across sessions, the original data satisfied a larger proportion of the predicted temporal criteria than did the corresponding shuffled data.

multiset, left/right counts, and evidence composition; the full workflow was repeated. The emphdecision-speed control kept units, choice labels, and both clustering stages fixed, permuted fast and slow labels within each choice condition while preserving group sizes, and rerun only Stage 2.

Every original or shuffled result was evaluated against six model-predicted temporal criteria. Two compound criteria required the summed activity of the selected action channel to exceed that of the opposite channel in all three early, middle, and late windows, separately for left- and right-choice trials. Four additional criteria tested the late-window pathway ordering: dL>iL and iR>dR on left-choice trials, and dR>iR and iL>dL on right-choice trials. The score was the percentage of these six criteria satisfied. For each session and control, the bar in Fig. S5 is the mean score over 50 repetitions. The 95% confidence interval is the percentile interval from 10,000 bootstrap resamples of the 50 scores. The unshuffled result was evaluated once and is shown as a black diamond.

### Post-Clustering Assessment of Potential FSI Inclusion

This assessment was applied after the four putative SPN labels had been assigned. For each assigned unit, we selected the mean extracellular waveform on the channel with the largest waveform amplitude. The waveform was baseline corrected, normalized to its maximum absolute amplitude, and aligned to its maximum negative deflection (Fig. S6A and Fig. S7A). Width at half of that negative deflection was measured from linearly interpolated crossings. Steinmetz waveforms used the native 30-kHz sampling rate; IBL waveforms came from the peak channel of each released array (82 samples $\times$ 32 channels). Whole-session firing rate was calculated from all spikes assigned to the unit. A unit was flagged as a putative FSI when both waveform width $< 0.15$ ms (Fig. S6B and Fig. S7B) and whole-session firing rate $> 10$ Hz (11, 13, 14). This joint criterion flagged 9 of 164 Steinmetz units (Fig. S6C) and 21 of 180 IBL units (Fig. S7C). The remaining sets contained 155 (94.5%) and 159 (88.3%) putative SPNs, respectively.

To assess whether putative FSIs affected inferred SPN subtype activity, we compared each subtype's 12-value pre-decision firing rate profile (six bins per choice) before and after excluding the flagged units. Pooled Pearson correlations were 0.998 for Steinmetz (Fig. S6D) and 0.997 for IBL (Fig. S7D); subtype-specific correlations were 0.992–1.000 and 0.996–0.999, respectively. Subsequent CLAW analyses excluded the 9 Steinmetz and 21 IBL units flagged as putative FSIs while retaining all other units with the inferred subtype labels.

### Construction of Striatal CLAWs

Based on our previously established CLAW (Circuit Logic Assessed via Walks) framework (2, 12), we represented continuous striatal activity as trajectories through a discrete state space. At each 10-ms time bin, striatal activity was encoded as a

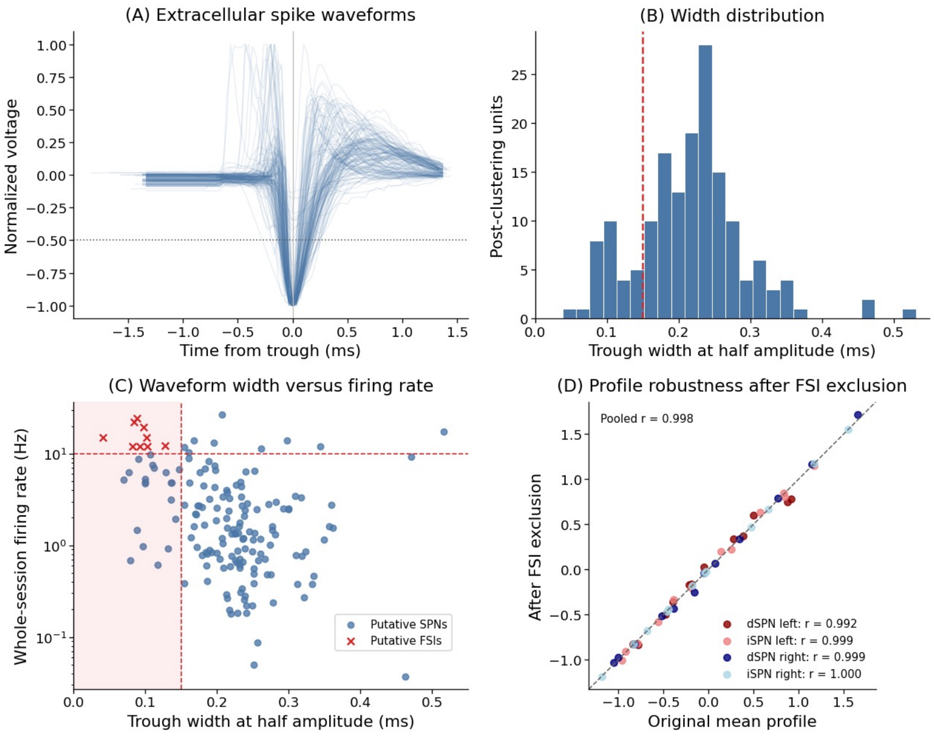


**Fig. S6.** Post-clustering assessment of potential FSI inclusion in the Steinmetz dataset. (A) Mean extracellular spike waveform for each of the 164 clustered units, measured on the channel with the largest waveform amplitude, aligned to the maximum negative deflection and normalized to its maximum absolute amplitude. The horizontal dotted line marks half of the maximum negative deflection used to measure waveform width. (B) Distribution of waveform widths. The red dashed line indicates the 0.15-ms threshold. (C) Joint classification based on waveform width and whole-session firing rate. Units were flagged as putative FSI when their waveform width was below 0.15 ms and their firing rate exceeded 10 Hz (11). Nine units met both criteria (red crosses); the remaining 155 units were retained as putative SPNs (blue circles). (D) Robustness of the subtype-averaged firing rate patterns to FSI exclusion. For each subtype, the pattern comprised 12 values: 6 time bins from left-choice trials and 6 from right-choice trials. Each point compares one value before and after exclusion of the putative FSI units; the dashed line indicates equality. The firing rate patterns were nearly unchanged after exclusion (pooled Pearson's $r = 0.998$; subtype-specific $r = 0.992$–$1.000$).

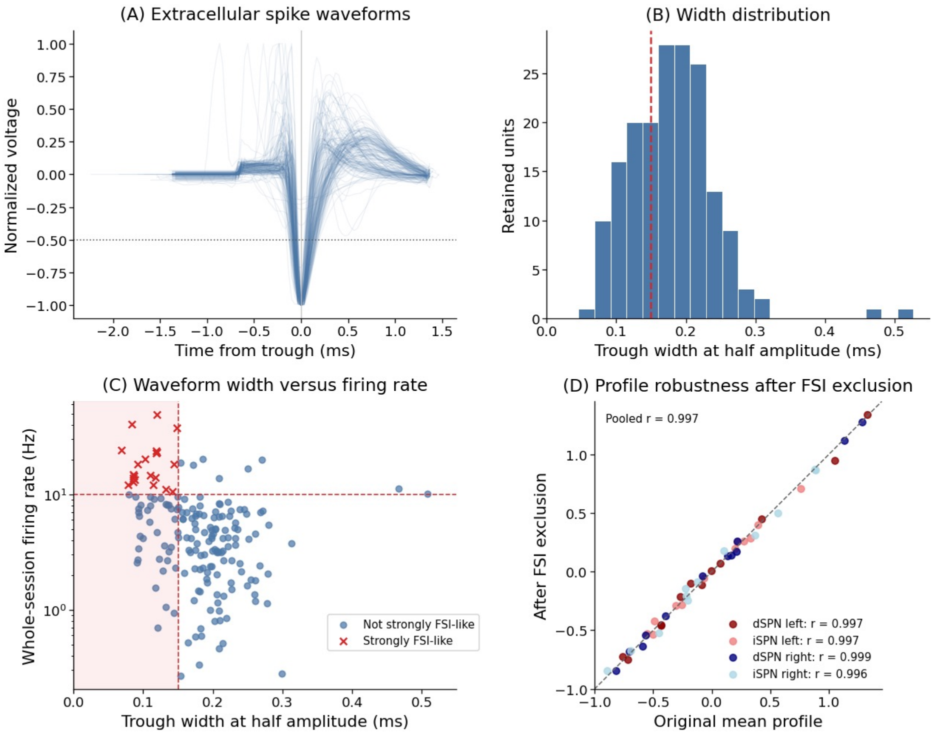


**Fig. S7.** Post-clustering assessment of potential FSI inclusion in the IBL dataset. Panels and notation are as in Fig. S6. Twenty-one units were flagged as putative FSIs (red crosses in panel C); the remaining 159 units were retained as putative SPNs. After FSI exclusion, the subtype-averaged pre-decision firing-rate patterns remained nearly unchanged (pooled Pearson's $r = 0.997$; subtype-specific $r = 0.996$–$0.999$).

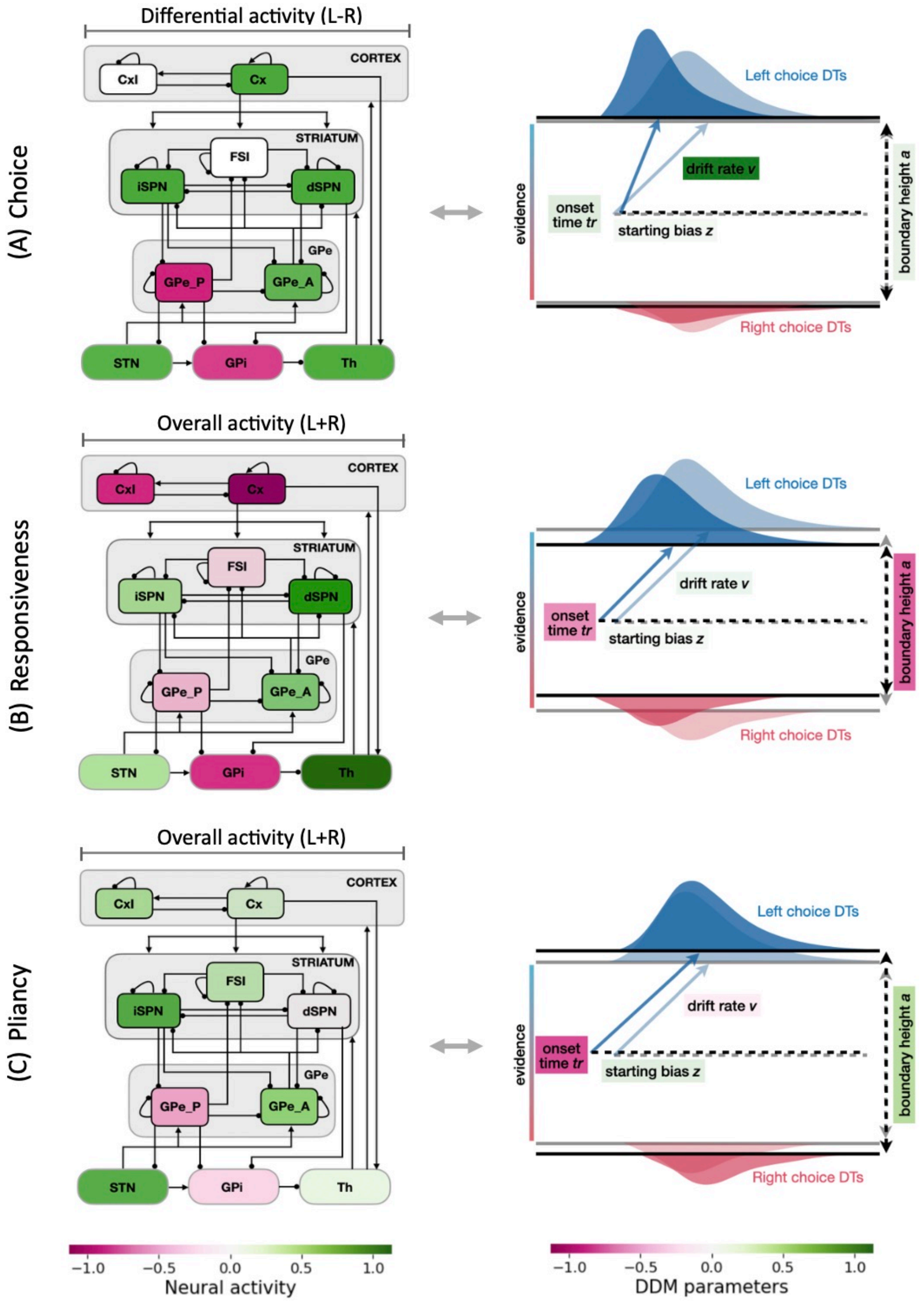


**Fig. S8.** Control ensembles and DDM correlates identified by canonical correlation analysis (CCA). Panels A, B, and C represent the relation between the firing rate of each population in the CBGT circuit and the DDM parameters obtained in the first three canonical components, labeled as choice, responsiveness, and pliancy, respectively. ach left subpanel represents neural contributions: for the choice ensemble (panel A), this is the differential activity between the left and right channels (L–R); for the responsiveness and pliancy ensembles (panels B and C), this is the overall activity summed across both channels (L+R). The other aspect of activity for each ensemble (L+R for choice and L–R for responsiveness/pliancy) is not shown, as it exhibits minimal impact on DDM parameters. Each right subpanel depicts the associated changes in the DDM parameters, as well as the resulting effect on the DT distributions. Figure reproduced from (12).

binary vector $s_k = [\mathrm{dL}, \mathrm{dR}, \mathrm{iL}, \mathrm{iR}]$, where each element indicated whether the corresponding SPN group was active or not. Each element was assigned a value of 1 when the corresponding population firing rate exceeded its binarization threshold and 0 otherwise. Following our prior analysis, the threshold for each population was set to the 90th percentile when its firing rate distribution was unimodal (Fig. S1D). For empirical SPN populations, thresholds were estimated separately within each session from their own firing rate distributions. Each binary vector was mapped to its base-2 integer representation, yielding 16 possible states numbered 0 through 15. Consecutive repetitions of the same state were compressed to one visit. Adjacent distinct states defined transitions, and the last pre-decision state was followed by END. For state $i$, $P(i \to j)$ was the number of $i$-to-$j$ events divided by all outgoing events from $i$, including END. Terminal probability was $P(\mathrm{END}|i)$ (labeled $P(\mathrm{end})$ in Fig. 3 in the main text). Left-choice probability (node color) and decision time (gray bar) were calculated across trials that

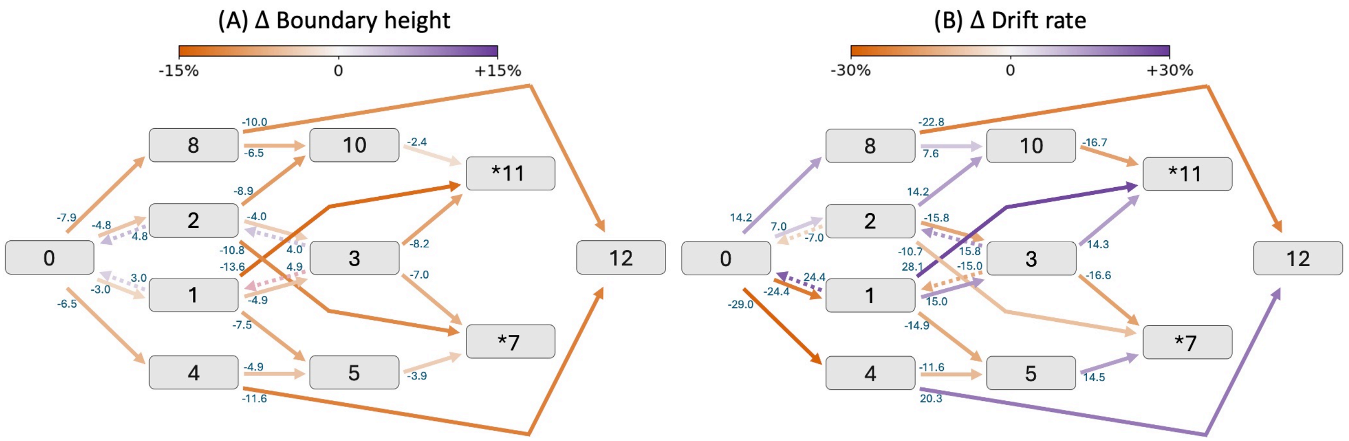


**Fig. S9.** Modulation of decision policy parameters by CBGT activity along the CLAW. The arrows indicate changes of the individual DDM parameters (boundary height, panel A; drift rate, panel B) associated with transitions between states of the CLAW. Changes were computed from the difference in the 18-dimensional firing rate activity between each state and the immediately downstream state. The magnitude of each change is indicated by the coloring (see color bar) and the number at the start of each arrow, representing the percentage change in parameter values.

visited each state; low-frequency edges were omitted only from the visualization. See (2) for complete details of the CLAW framework and analysis.

### Statistical Analysis of iSPN Roles

We tested the three state-dependent predictions illustrated in Fig. 4 of the main text. Model and empirical data were analyzed separately, with eligible trials pooled across networks or sessions. Uncertainty was assessed by comparing left-choice probability among trials visiting iSPN-only states 1, 2, or 3, left-channel dSPN states 8 or 10, and right-channel dSPN states 4 or 5. A trial could contribute to multiple groups if its trajectory visited more than one category. Two contrasts were calculated in each bootstrap replicate: iSPN-only versus left-channel dSPN states and iSPN-only versus right-channel dSPN states. In each of 5,000 bootstrap iterations, trials were sampled with replacement from each group using the size of the smallest group as the common sample size, and left-choice probability was calculated. The resulting bootstrap distributions were compared by one-way ANOVA, followed by two-sided $t$ tests comparing the iSPN-only group with each dSPN group, with Bonferroni correction for the two pairwise comparisons.

Commitment was assessed by comparing terminal probability for state 8 with that for state 10 entered directly from state 8, and for state 4 with that for state 5 entered directly from state 4. Trials visiting the corresponding source state were resampled, and each pair of bootstrap distributions was compared using a two-sided $t$ test. The decision-slowing analysis used raw decision time distributions without bootstrap resampling. Along the left-biased path, trials visiting state 10 without a later visit to state 11 were compared with trials in which state 11 occurred later in the sequence. The corresponding right-path comparison used state 5 without a later visit to state 7 versus state 5 followed later by state 7. Two-sided $t$ tests were performed separately for each path.

### Control Ensembles and DDM Projection

The DDM parameters and the CBGT activity for our 300 network configurations were used to identify control ensembles through canonical correlation analysis (CCA), as was done in prior work (2, 3, 15). For each of the 300 simulated networks, distributions of choice and decision time were fit with a static drift-diffusion model using the Hierarchical Sequential Sampling Modeling (HSSM) toolbox in Python, yielding the corresponding configuration of static DDM parameters (boundary height $a$, drift rate $v$, onset time $tr$, and starting bias $z$) for each network. CCA was then applied to relate these parameters to an 18-dimensional representation of CBGT activity, consisting of the 2 shared interneuron activities, 8 channel-summed activity measures, and 8 between-channel differences. The resulting canonical variates defined correlated patterns of CBGT activity and DDM parameters, and the three leading components defined the choice, responsiveness, and pliancy ensembles (Fig. S8).

To map striatal CLAW transitions onto control ensembles, we used the full 18-dimensional CBGT activity vector from each simulated network. For each network and striatal state, activity was averaged across all pre-decision bins assigned to that state. A transition from source state $i$ to target state $j$ was evaluated in networks in which both states occurred, and the corresponding activity change was calculated as $\Delta F_{ij} = F_j - F_i$. Changes in neural canonical scores were obtained as $W_{ij} = \Delta F_{ij} U$, where $U$ contains the neural CCA loadings. Associated changes in DDM parameter space was calculated as $W_{ij} V^T$, where $V$ contains the DDM loadings (Fig. S9). Transition summaries were calculated as the median across networks containing both states. This analysis was restricted to the CBGT CLAW because the empirical recordings lacked simultaneous activity and identities for the full CBGT population set.

### References


1. M Clapp, et al., Cbgtpy: An extensible cortico-basal ganglia-thalamic framework for modeling biological decision making. *PLoS One* **20**, e0310367 (2025).
2. Z Yu, T Verstynen, JE Rubin, How the dynamic interplay of cortico-basal ganglia-thalamic pathways shapes the time course of deliberation and commitment. *PLOS Comput. Biol.* **22**, e1012966 (2026).
3. J Bahuguna, T Verstynen, JE Rubin, How cortico-basal ganglia-thalamic subnetworks can shift decision policies to increase reward rate. *PLOS Comput. Biol.* **21**, e1013712 (2025).
4. NA Steinmetz, P Zatka-Haas, M Carandini, KD Harris, Distributed coding of choice, action and engagement across the mouse brain. *Nature* **576**, 266–273 (2019).
5. International Brain Laboratory, et al., A brain-wide map of neural activity during complex behaviour. *Nature* **645**, 177–191 (2025).
6. S Amadio, et al., Motor evoked potentials in a mouse model of chronic multiple sclerosis. *Muscle & Nerve: Off. J. Am. Assoc. Electrodiagn. Medicine* **33**, 265–273 (2006).
7. V Castoldi, E Rossi, S Marenna, G Comi, L Leocani, Improving reproducibility of motor evoked potentials in mice. *J. Neurosci. Methods* **367**, 109444 (2022).
8. JH Ward Jr, Hierarchical grouping to optimize an objective function. *J. Am. statistical association* **58**, 236–244 (1963).
9. HW Kuhn, The hungarian method for the assignment problem. *Nav. research logistics quarterly* **2**, 83–97 (1955).
10. GN Lance, WT Williams, A general theory of classificatory sorting strategies: 1. hierarchical systems. *Comput. J.* **9**, 373–380 (1967).
11. JM Burkhardt, X Jin, RM Costa, Dissociable effects of dopamine on neuronal firing rate and synchrony in the dorsal striatum. *Front. integrative neuroscience* **3**, 562 (2009).
12. Z Yu, JE Rubin, T Verstynen, Learning reweights the decision dynamics of cortico-basal ganglia-thalamic pathways from deliberation to commitment. *bioRxiv* (2026).
13. GJ Gage, CR Stoetzner, AB Wiltschko, JD Berke, Selective activation of striatal fast-spiking interneurons during choice execution. *Neuron* **67**, 466–479 (2010).
14. JD Berke, Uncoordinated firing rate changes of striatal fast-spiking interneurons during behavioral task performance. *The J. Neurosci.* **28**, 10075–10080 (2008).
15. C Vich, M Clapp, JE Rubin, T Verstynen, Identifying control ensembles for information processing within the cortico-basal ganglia-thalamic circuit. *PLOS Comput. Biol.* **18**, e1010255 (2022).
